# Self-renewing Sox9+ osteochondral stem cells in the postnatal skeleton

**DOI:** 10.1101/2023.12.07.570646

**Authors:** Stephanie Farhat, Bahaeddine Tilouche, Spencer Short, Medjie Piron, T. Mark Campbell, Alex Fernandes, Mariya Somyk, Hina Bandukwala, Eric Arezza, Quentin Sastourne-Arrey, Katherine Reilly, Maria Abou Chakra, Gary Bader, Leo Kunz, Timm Schroeder, Sasha Carsen, Pierre Mattar, Jeffrey Dilworth, Daniel L. Coutu

## Abstract

Postnatal skeletal growth, homeostatic maintenance, and regeneration is driven by skeletal stem cells. In addition, it is well established that skeletal tissues lose their regenerative potential with age, comorbidities, and repeated trauma, possibly through stem cell exhaustion or loss of function. However, it is largely unknown where these cells reside in skeletal tissues, what molecular mechanisms regulate their self-renewal and fate decisions, and how to isolate, purify, and expand them ex vivo. Therefore, there is an urgent need for a deeper understanding of postnatal skeletal stem cells. Here, we used genetic lineage tracing, thymidine analogues retention, whole bone microscopy, imaging cytometry, in vitro assays, and single cell transcriptomics and provide the first experimental evidence for the existence of self-renewing osteochondral stem cells in the postnatal skeleton in both males and females. We also show direct comparisons between adult, fetal, mouse, and human skeletal stem cells at the transcriptome level.

---

Skeletal trauma, degeneration, or congenital diseases affect over 50% of the population worldwide at least once in their lifespan. In young individuals, skeletal tissues can regenerate almost seamlessly. However, this regenerative capacity gradually decreases with age, comorbidities, and repeated trauma ^1–6^. Similarly, surgery and/or rehabilitation aimed at preserving, reconstructing, or replacing injured tissues have a good success rate in younger individuals but often fail with increasing age. Musculoskeletal diseases now constitute the most common reason people visit their physicians, making up the world’s most common chronic diseases. Recent experimental approaches to enhance skeletal tissue repair involve stimulation of endogenous repair mechanisms, cell transplantation, and tissue engineering. Yet, none of these techniques have achieved standard clinical use due to lack of efficacy. With the growing global burden of musculoskeletal diseases, there is therefore a need to better study skeletal stem cells (SSCs) in order to develop novel stem cell-based regenerative therapies in orthopedics.

Skeletal tissues, like all somatic tissues with a high cell turnover, rely on stem cells for their postnatal development, homeostatic maintenance, and regeneration after injury. However, the cellular phenotype, molecular characteristics, and anatomical niche of SSCs remains elusive to date. Several attempts have been made to identify SSCs using cell surface markers^7–16^, reporter mouse lines^17,18^, and genetic lineage tracing^7,19–31^. These studies focused mainly on embryonic or perinatal developmental stages and provide important information about the ontogeny and relationship between various skeletal cell lineages. More recently, studies by various groups suggested that some cells residing in the resting zone of the physeal cartilage (growth plate) could act as stem cells driving postnatal longitudinal bone growth^32^. However, none of these studies were able to definitively demonstrate multipotency and self-renewal of these putative SSCs.

The existence and location of postnatal SSCs has been difficult to study using lineage tracing mainly because of technical challenges; indeed, bone tissue is very hard and difficult to process using normal histological techniques and is also highly autofluorescent. We recently developed techniques addressing these hurdles as well as computational methods to perform 3D imaging cytometry on large postnatal bones^33–35^. Using these techniques, we here show that a subpopulation of cells expresses high levels of Sox9, possesses osteochondral properties in vivo and in vitro, self-renews long-term in growing and mature bones, and can be isolated and expanded ex vivo. These cells are abundant in the resting zone of the physes and are also found in periosteum. Single cell transcriptomics provides information about the molecular characteristics of these SSCs, and we show that postnatal human skeletal tissues also harbor cells with similar features.

## Results

### Resting zone Sox9+ cells are multipotent and self-renewing in postnatal bones

Recent studies suggest that putative SSCs in the physis express chondrocyte markers as well as genes involved in early embryonic skeletal development. Since the transcription factor Sox9 is one of the earliest genes expressed during the development of the appendicular skeleton and is a master regulator of chondrogenesis^36^, we obtained Sox9-CreERT transgenic mice and crossed them with the Ai14 Cre-reporter line (R26^tdTomato^). In the resulting line, tamoxifen (TAM) administration leads to constitutive expression of tdTomato (tdT) in Sox9+ cells and their progeny. We performed dual lineage tracing by pulsing 8-week-old (postnatal day 56, P56) animals with TAM and the thymidine analogue EdU for three consecutive days, and retrieved femurs at two, 30 and 90 days post-labeling (Fig.1A). This allowed us to track the fate of Sox9+ cells and their progeny based on tdT expression, to quantify proliferative cells and track their fate based on EdU incorporation, and to measure cell turnover rate based on tdT and EdU label retention. Harvested femurs were fixed, decalcified, sectioned (300-500μm thickness, representing 30-50% of the entire bone volume), stained (Sox9 antibody and Click-IT EdU detection), optically cleared, and the entire sections were scanned using spectral confocal imaging. We next performed 3D segmentation to identify tdT+ cells in the resulting images, and quantified fluorescence intensity of Sox9 and EdU within tdT+ cells using our open-source software XiT^35^. Using XiT, Sox9 protein expression and EdU incorporation by tdT+ cells can be quantified based on staining controls (Fig.S1) and displayed as scatter plots or spatial distributions (Fig.1B). Quality control is performed by selecting tdT+ cells that co-express Sox9 and incorporated EdU, exporting these cells’ identifiers, and visually confirming that these segmented objects represent tdT+Sox9+EdU+ cells in the original images (Fig.1C).

**Figure 1.**
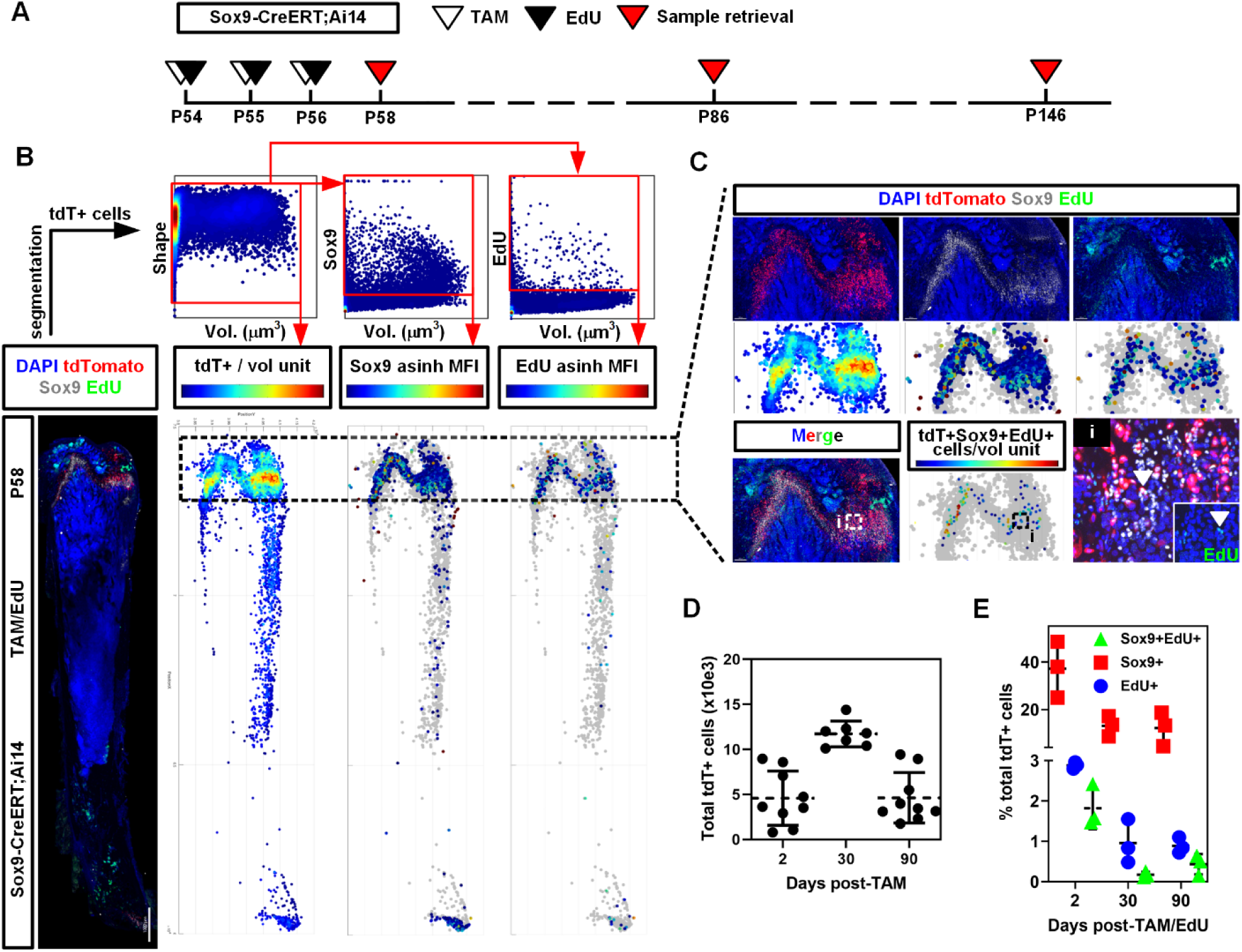
Identification of self-renewing skeletal stem cells using dual lineage tracing. A) Sox9-CreERT;Ai14 mice were pulsed with TAM and EdU injections for three consecutive days (P54, P55, P56) and femurs were collected after chases of two, 30, and 90 days. B) Spectral confocal imaging of optically-cleared, 300μm-thick whole femur sections is performed and tdT+ cells are identified using image segmentation. Histomorphometric data of tdT+ cells is derived along with their expression in Sox9 protein (based on antibody staining) and EdU incorporation, and displayed as scatter plots or spatial (anatomical) representation. Arcsinh (asinh) of the mean fluorescence intensity (MFI) is used to better visualize small and large difference in MFI. C) Quality control is performed by ensuring that the cell phenotypes observed by thresholding (based on staining controls) matches the observed phenotypes in the original images. D) Quantification of total tdT+ cells detected per section in each of the three lineage traces (days 2, 30 and 90 post-labeling, n=9 sections from three males and four females for each trace). E) Quantification of Sox9+, EdU+, and Sox9+EdU+ cells in the tdT+ populations, expressed as the percentage of total tdT+ cells (individual values are shown as well as mean +/-SD. n=3 (males only) for each data point.

We detected an average of 4.58+/-1.50×10^3^ tdT+ cells per section (n=9) at day two post-labeling (Fig.1D). Of these, 37.2+/-5.5% had detectable Sox9 protein, 2.89+/-0.04% had incorporated EdU, and 1.82+/-0.26% were tdT+Sox9+EdU+ (Fig.1E, n=3 for each lineage trace). During bone growth, the number of tdT+ cells increased to 11.7+/-0.7×10^3^. When bones entered the homeostatic maintenance phase and stopped growing, the number of tdT+ cells stabilized to that observed in short chases, i.e. 4.6+/-1.4×10^3^. The fraction of tdT+Sox9+ also stabilized over time to 13.1+/-2.1% and 12.2+/-3.6%, 30- and 90-days post-labeling, respectively. However, the fraction of tdT+EdU+ and tdT+Sox9+EdU+ steadily decreased over time, suggesting that Sox9+ cells proliferate slowly but continuously in postnatal bones.

At day two post-labeling we observed abundant tdT+ chondrogenic cells in the physes of the femur (Fig.2A). We also observed tdT+ osteogenic cells in trabecular and cortical bone, as well as periosteum. These cells were more abundant close to the physes. tdT+ cells with detectable Sox9 protein and/or EdU incorporation were found mainly in the resting zone of the physes (Fig.1B-C, 2A) but also in periosteum and cortical bone, where their abundance was negatively correlated with their distance from the physes (Fig.1B). In the physes, most of the tdT+ cells were found in the resting zone and appeared as small and round, Sox9+ cells. Only a few resting zone cells had incorporated EdU during cell division (Figs.1C, 2A). Columns of proliferating chondrocytes were largely EdU+ but tdT-, despite showing detectable levels of Sox9 protein (see Discussion below). In contrast, most tdT+ osteogenic cells in metaphyseal trabecular and cortical bone were Sox9-EdU- (Fig.2A). We also observed abundant tdT-EdU+ cells in the periosteum, throughout the metaphysis and diaphysis (Fig.2A).

**Figure 2.**
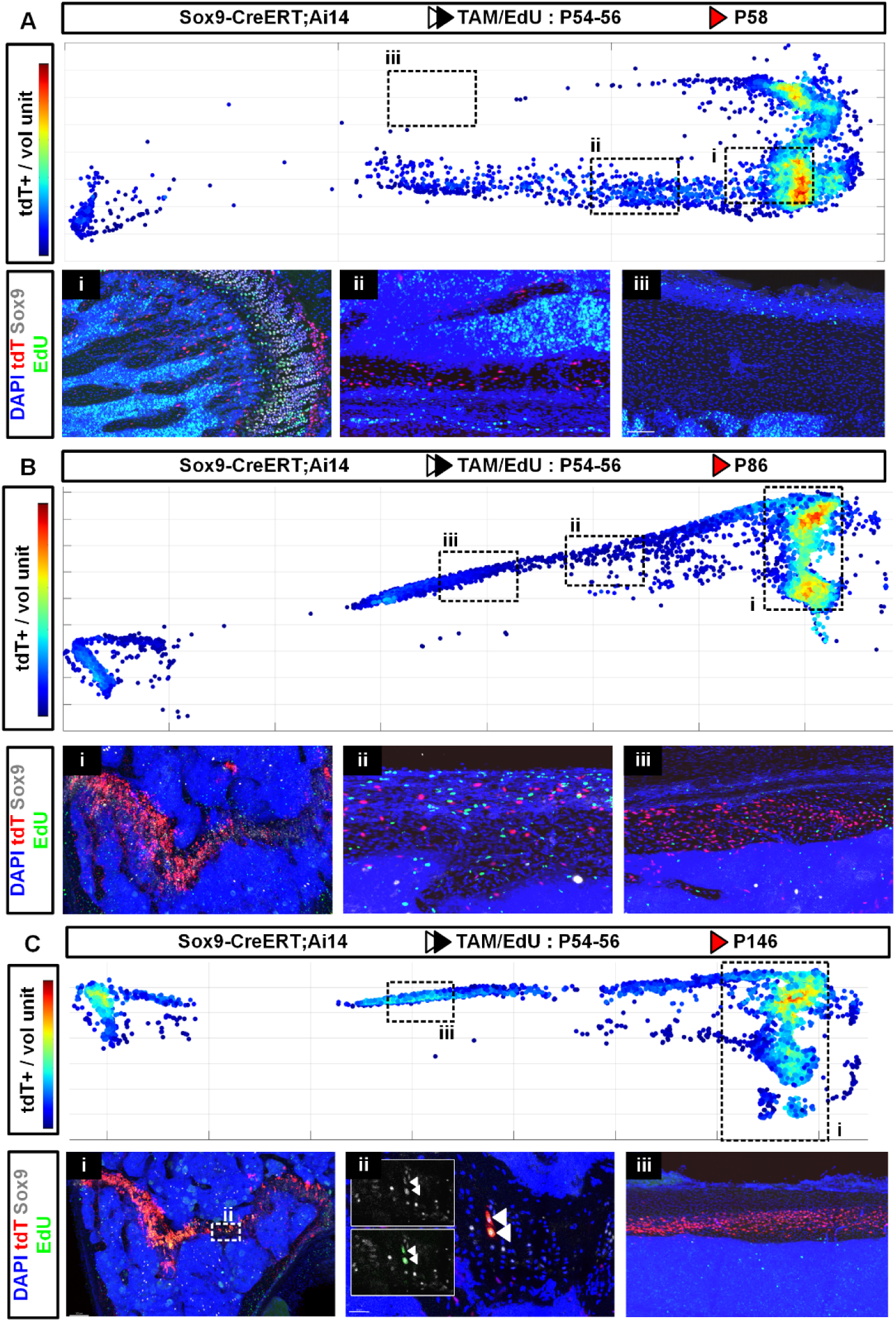
Dual lineage tracing of Sox9+ and proliferative cells in the postnatal mouse skeleton. Sox9-CreERT;Ai14 mice received TAM and EdU on three consecutive days (P54-P56), femurs were retrieved on days 30 and 90 post-labeling to assess the fate of Sox9+ and proliferative cells. A) Anatomical distribution of tdT+, Sox9+ and EdU+ cells, two days post-labeling. B) Anatomical distribution of tdT+, Sox9+ and EdU+ cells, 30 days post-labeling. C) Anatomical distribution of tdT+, Sox9+ and EdU+ cells, 90 days post-labeling. Experiment was performed in three female and three male mice for each trace. Representative images are shown (male mice only).

At day 30 post-labeling, most of the chondrogenic cells in the physes were tdT+ (Fig.2B) and we also observed an increase in the number of tdT+ osteogenic cells in trabecular and cortical bone. Only a few rare cells retained EdU in the physes and these were largely tdT+ (Fig.S2). tdT-EdU+ cells were still detectable in the metaphysis and diaphysis, but were now found deeply embedded in cortical bone, suggesting differentiation into post-mitotic osteocytes. This further indicates rapid osteogenic cell turnover in growing bones, with a large fraction of osteocytes being replaced by periosteal progenitors within 30 days. At day 90 post-labeling, the mice were skeletally mature and their bones had stopped growing. In the physes, columns of proliferating chondrocytes were no longer visible but tdT+ cells were still abundant (Fig.2C). Remarkably, rare tdT+Sox9+EdU+ cells were still observed in the resting zone, often appearing as doublets of cells that had just divided (Fig.2C,ii, white arrowheads). In cortical bone, no EdU+ cells were observed but tdT+ cells remained abundant (Fig.2C,iii). In 30- and 90-day chases, we also observed the appearance of tdT+ osteogenic cells in the epiphysis and subchondral bone (Fig.S3), structures that were tdT-in shorter chases. In the periosteum, we could not detect tdT+Sox9+ cells in 90-day chases, indicating that either periosteal Sox9+ cells do not self-renew, or their abundance falls below our detection threshold in older animals.

Taken together, these results indicate that Sox9-CreERT is expressed by both chondrogenic and osteogenic cells in postnatal bones. Most of these cells are highly proliferative to sustain bone growth and elongation, as well as rapid skeletal tissue turnover. However, a subpopulation of Sox9+ cells residing in the resting zone of the physes is slow-cycling and appears to give rise to both osteogenic and chondrogenic cells in the metaphysis, diaphysis, and epiphysis. Some of these small and round cells persist for months in the resting zone until skeletal maturity, maintain expression of both tdT and Sox9 protein, and retain proliferation capacity (Fig.S4). Hence, these display features characteristic of self-renewing osteochondral stem cells. Intriguingly, we observed tdT+ adipocytes in the bone marrow of female mice but never in the bone marrow of males (Fig.S5).

### Self-renewing osteochondral Sox9+ SSCs persist in the mature skeleton

To confirm that Sox9+ SSCs persist in the mature skeleton and remain multipotent and self-renewing, we performed longer chases (up to six months – 180 days) after labeling Sox9+ cells in young (8-week-old, P56) and skeletally mature (P180) mice. Moreover, since Sox9 and its inducible tdT Cre-reporter were observed in various cells of the chondrocyte and osteocyte lineages, we sought to simplify the interpretation of the lineage tracing data by labeling only the cells in which the Sox9 Cre-reporter transgene was being highly transcribed. For this, we pulsed the mice with a single injection of 4-hydroxytamoxifen (4OHT), an active metabolite of TAM that is only bioavailable for two hours in mice^37^. Since skeletal physiology exhibits sexual dimorphism, mainly due to the effects of estrogens on skeletal cells, we also analyzed female and male mice separately.

When Sox9+ cells were labeled in P56 mice, the anatomical distribution of tdT+ cells over time was similar to that observed in longer pulses (Fig.3A and Fig.S6, compare with Fig.1-2). We observed minor but statistically significant differences between young females and males (Fig.3B and C, left panels), suggesting that female bones undergo an accelerated postnatal development compared to males. These differences were attenuated in older animals (Fig.3B and C, right panels). In younger animals, the total number of tdT+ cells initially increased during bone growth, then stabilized in mature bones (Fig.3B). The number of tdT+Sox9+ cells initially decreased before stabilizing in more mature bones, reflecting the decreased number of Sox9+ proliferating chondrocytes in the growth plate. In young animals, we observed a 40:60 chondrocyte:osteocyte ratio in tdT+ cells across timepoints, whereas in older animals the fate of tdT+ cells appeared slightly biased towards the chondrogenic fate in what remained of the physes (Fig.3C).

**Figure 3.**
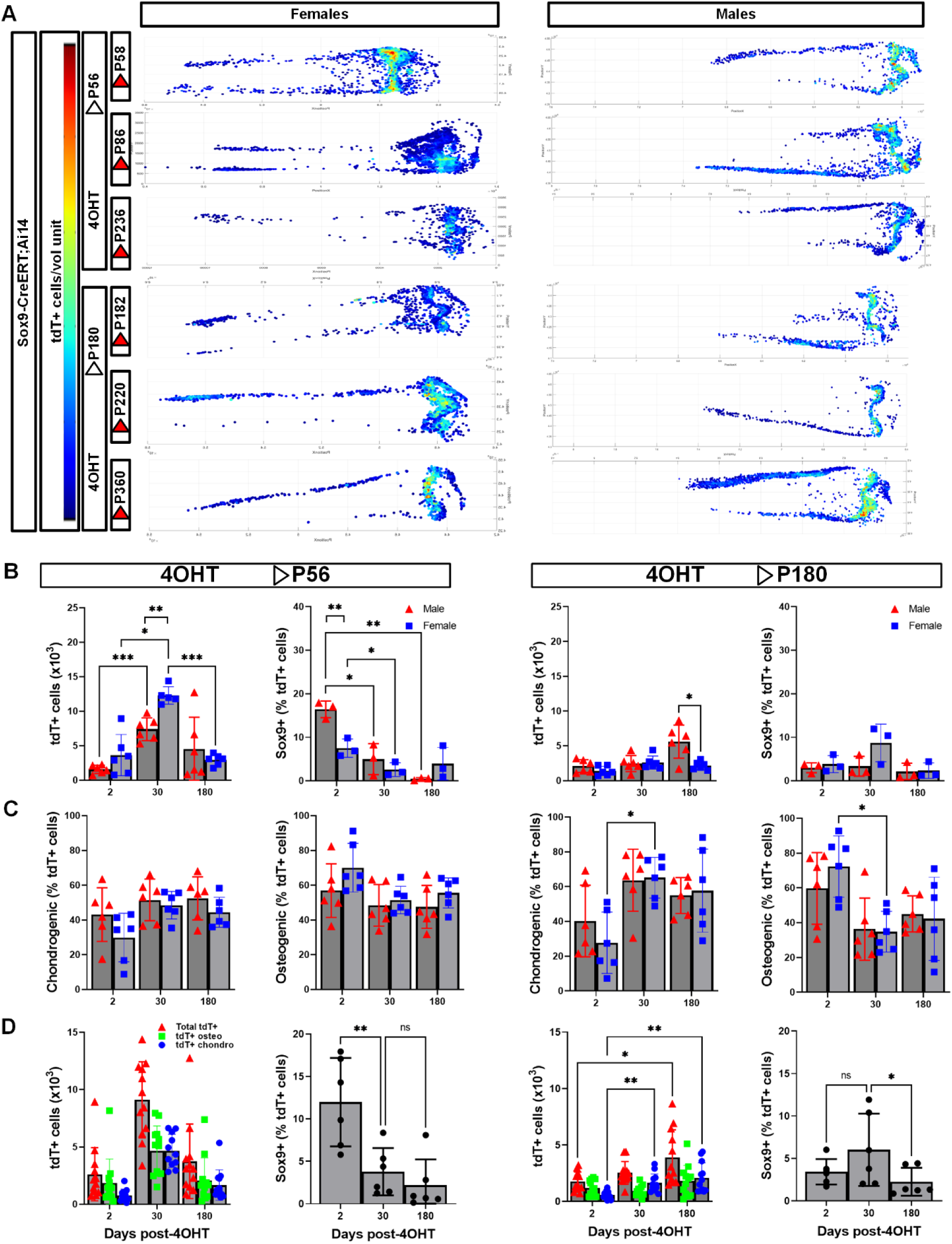
Persistence of self-renewing Sox9+ SSCs in skeletally mature male and female mouse bones. Sox9-CreERT;Ai14 mice were pulsed with a single injection of 4OHT at P56 or P180 and chase for up to 180 days post-labeling. A) Anatomical distribution of tdT+ cells over time when P56 and P180 female and male mice were pulsed. B) Quantification of total tdT+ cells (n=6) and tdT+Sox9+ cells (n=3) in different lineage traces, with comparison between female and male mice. C) Quantification of tdT+ chondrogenic and osteogenic cells over time in female and male mice (n=6 for both females and males) in each trace. D) Pooled data from female and male mice showing the total number of tdT+ cells, tdT+ chondrogenic cells, tdT+ osteogenic cells (n=12 each), and percentage of tdT+Sox9+ cells (n=6 each) over time in different lineage traces.

When juvenile female and male data are pooled, the number of cells initially labeled was lower (2.6+/-1.2×10^3^ cells/section, n=12) but reached similar levels as with longer pulses after 30 days (9.1+/-3.2×10^3^) (Fig.3D, left). In 180-day chases, the number of tdT+ cells was also similar to that observed in 90-day chases with longer pulses (3.8+/-1.7×10^3^). The number of tdT+ chondrogenic and osteogenic cells followed a similar kinetics over time. We however observed a sharper decrease in the fraction of tdT+Sox9+ cells (presumably self-renewing SSCs) from 12.9+/-3.3% of total tdT+ cells at day two post-labeling, to 2.2+/-1.5% at day 180 (Fig.3D, n=6 each).

Similarly, when we pooled male and female data from skeletally mature mice having received a short 4OHT pulse at P180, the anatomical distribution and number of tdT+ cells in short, two-day chases was strikingly similar to that observed in younger animals (Fig.3D, right), with 1.8+/- 0.4×10^3^ tdT+ cells observed per section (n=12). However, instead of a biphasic population kinetics (increase in cell number followed by a decrease) we observed a gradual increase in the number of tdT+ cells, reaching 3.9+/-1.2×10^3^ by day 180 (Fig.3D, right), which is again similar to the number of cells observed in younger animals after 180-day chases. The fraction of tdT+Sox9+ cells observed was highly variable in two-day chases but remained mostly constant in longer chases (6.0+/-2.2% and 3.5+/-1.2% after 30 and 180 days, respectively). In summary, our data indicates that self-renewing and multipotent Sox9+ SSCs persist in the skeleton throughout postnatal development and in mature bones.

### Sox9+ SSCs proliferate and retain multilineage differentiation potential in vitro

To assess if Sox9+ SSCs could be purified and expanded ex vivo for further characterization or transplantation assays, we performed enzymatic dissociation using collagenases and neutral protease followed by gentle mechanical dissociation on postnatal mouse femurs. We then used magnetic cell separation (negative selection) to remove contaminating endothelial and hematopoietic cells. Using this method, we typically achieve a cell viability >90% (Fig.4A, left panel). However, flow cytometry analysis demonstrated that <0.1% of these skeletal cells expressed tdT (Fig.4A, right panel). This is 10-fold lower than we expected based on the percentage of tdT+ cells in total skeletal cells we observed using imaging (Fig.4B).

**Figure 4.**
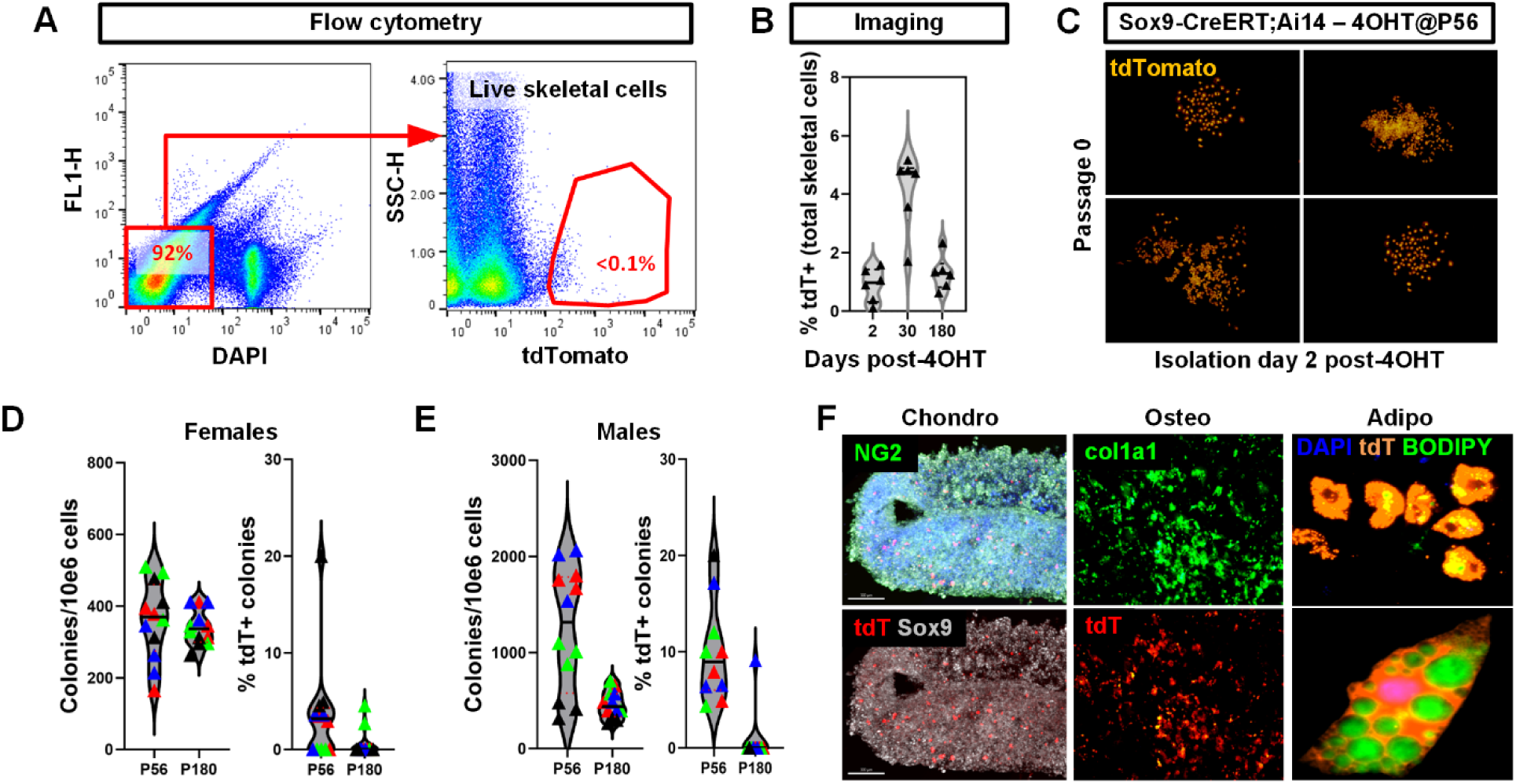
Isolation, culture expansion and in vitro multilineage differentiation of Sox9+ mSSCs. Sox9-CreERT;Ai14 mice were pulsed with a single 4OHT injection at P56 or P180, as indicated. Femurs were harvested two days post-labeling and skeletal cells isolated using enzymatic and mechanical dissociation, followed by magnetic removal of endothelial and hematopoietic cells through negative selection. A) Flow cytometry analysis of skeletal cell viability after isolation (left) and quantification of tdT+ cell number (right). B) Quantification of the number of tdT+ cells in femur sections expressed as the percentage of total skeletal cells, based on imaging data. The number of total skeletal cells was obtained by segmenting cell nuclei in trabecular and cortical bone, as well as the physes, but excluding cells in bone marrow. N=6 each. C) After isolation, tdT+ cells in vitro formed colonies of cells with varying morphologies and cell numbers. D, E) Quantification of the total number of colonies obtained per 10^6^ skeletal cells seeded and the fraction of these colonies that were tdT+, expressed as the percentage of total colonies. The analysis was performed in young (P56) and adult (P180) female and male mice (D and E, respectively). Shown are violin plots of the data distribution, overlayed with individual values and median values. N=12 for females and N=9 for males. The biological replicates are color coded, and each was performed in three separate technical replicates. F). Mixed populations of skeletal cells (tdT+ and tdT-) were culture expanded for three passages then subjected to differentiation into chondrocytes, osteoblasts, and adipocytes. Differentiation was assessed using iμμunostaining and the presence of differentiated tdT+ cells was qualitatively assessed. Representative images shown. The assay was performed on three biological replicates, each in three separate technical replicates.

When we pulsed P56 mice with 4OHT and purified tdT+ cells using FACS or cultured them in individual wells, they failed to form colonies. However, when unfractionated skeletal cells were seeded at clonal densities in large cell culture vessels, we observed the formation of several tdT+ colonies (Fig.4C). We then quantified the total number of colonies formed and the fraction of tdT+ colonies, in young and adult mice of each sex. In female mice, the total number of colonies obtained did not vary significantly between young and adult mice (360+/-54 and 341+/-25 colonies per 10^6^ skeletal cells, respectively, n=12) (Fig.4D). In young females, 3.6+/-2.8% of the colonies were tdT+, but this fraction decreased to 0.6+/-0.7% in adult females. In young male mice, we obtained significantly more colonies compared to females (1535+/-220 colonies per 10^6^ skeletal cells) (Fig.4E, n=9). This number decreased but remained elevated in adult males (517+/-108 colonies per 10^6^ skeletal cells). The fraction of tdT+ colonies was also higher in young males compared to females (8.8+/-2.0% of total colonies) but also decreased significantly in older males (1.0+/-1.5%). In summary, Sox9+ cells can be expanded in vitro for six to seven low density passages (split every ten days) but their specific growth requirements ex vivo, especially when isolated from older animals, remain to be determined.

We next tested if isolated tdT+ cells retained multilineage differentiation potential after culture-expansion. Mice were pulsed with 4OHT at P56 and skeletal cells were isolated from femurs 48h later. We monitored the heterogenous cultures for the presence of tdT+ colonies. Wells that contained tdT+ colonies were selected and further expanded for three passages to obtain sufficient cells for differentiation assays. The cells were placed in osteogenic or adipogenic media, or aggregated and placed in chondrogenic medium. Differentiation was assessed by immunostaining for specific markers and differentiated cells were assayed for their expression of tdT. Our results indicate that polyclonal colonies of tdT+ cells derived from Sox9+ cells maintain their multilineage differentiation potential in vitro after extensive proliferation and expansion (Fig.4F).

### Postnatal Sox9+ SSCs express genes involved in chondrogenesis, osteogenesis and self-renewal

To obtain a detailed molecular characterization of postnatal Sox9+ SSCs, we sought to use single cell transcriptomics data of postnatal skeletal cells. We took advantage of a recently published, open-access dataset of skeletal cells that contain mostly chondrogenic and osteogenic cells^38^. In this dataset, several clusters of cells expressed Sox9 (Fig.5A-B). Several of those also expressed genes specific to articular cartilage and joint tissues (Scx, Prg4, Tbx15, Tbx18) and were not analyzed further. We focused our analysis on clusters 2, 10, 13 and 17 (circled in Fig.5B). To identify which of these clusters contained self-renewing SSCs, we performed a gene length analysis of the transcripts expressed by the cells in those clusters. Indeed, fast cycling cells tend to express shorter genes and accumulate less transcripts per cell^39^. Slow cycling, quiescent stem cells should thus have time to transcribe longer gene and accumulate more transcripts. We found that cells in clusters 13 and 17 expressed significantly longer transcripts than those from clusters 2 and 10 (Fig.5C). We performed differential gene expression (DGE) analysis on these four clusters (Fig.5D, Supplemental Tables 1-4) and then a gene ontology (GO) term analysis on these differentially expressed genes. Cells from cluster 2 differed from the other clusters by the fact they downregulated most genes, hence GO term analysis was not possible. However, cells from cluster 10 expressed mostly genes involved in mitosis and cell division (Fig.5E). Cluster 13 cells expressed mostly genes involved in inhibition of Wnt signaling (e.g., Wif1, Sfrp5) as well as bone development, while cluster 17 cells also expressed gene mostly involved in bone and mesenchyme development (Fig.5E). Wif1 and Sfrp5 have already been described to be expressed and regulate resting zone chondrocytes^40^, and we found that they were both highly specific to cells in cluster 13 (Fig.5F, Sfrp5 shown). We confirmed that Sfrp5 was specifically expressed in resting zone chondrocytes using immunostaining (Fig.5G). In order to better understand the signaling pathways that regulate Sox9+ SSCs self-renewal and where these signals originated from, we next perform a CellChat analysis^41^. We found that chondrocytes, osteoblast lineage cells, and perichondral cells were important sources of canonical and non-canonical Wnt signalling to the resting zone chondrocytes, whereas only perichondral cells produced Hh signals (Fig.5H). On the other hand, cells described as fibroblastic-like^38^ but likely of articular cartilage or joint tissue origin (based on gene expression) where important sources of FGF and PTH signals. Finally, we found that resting zone chondrocytes themselves were an important source of PDGF signals.

**Figure 5.**
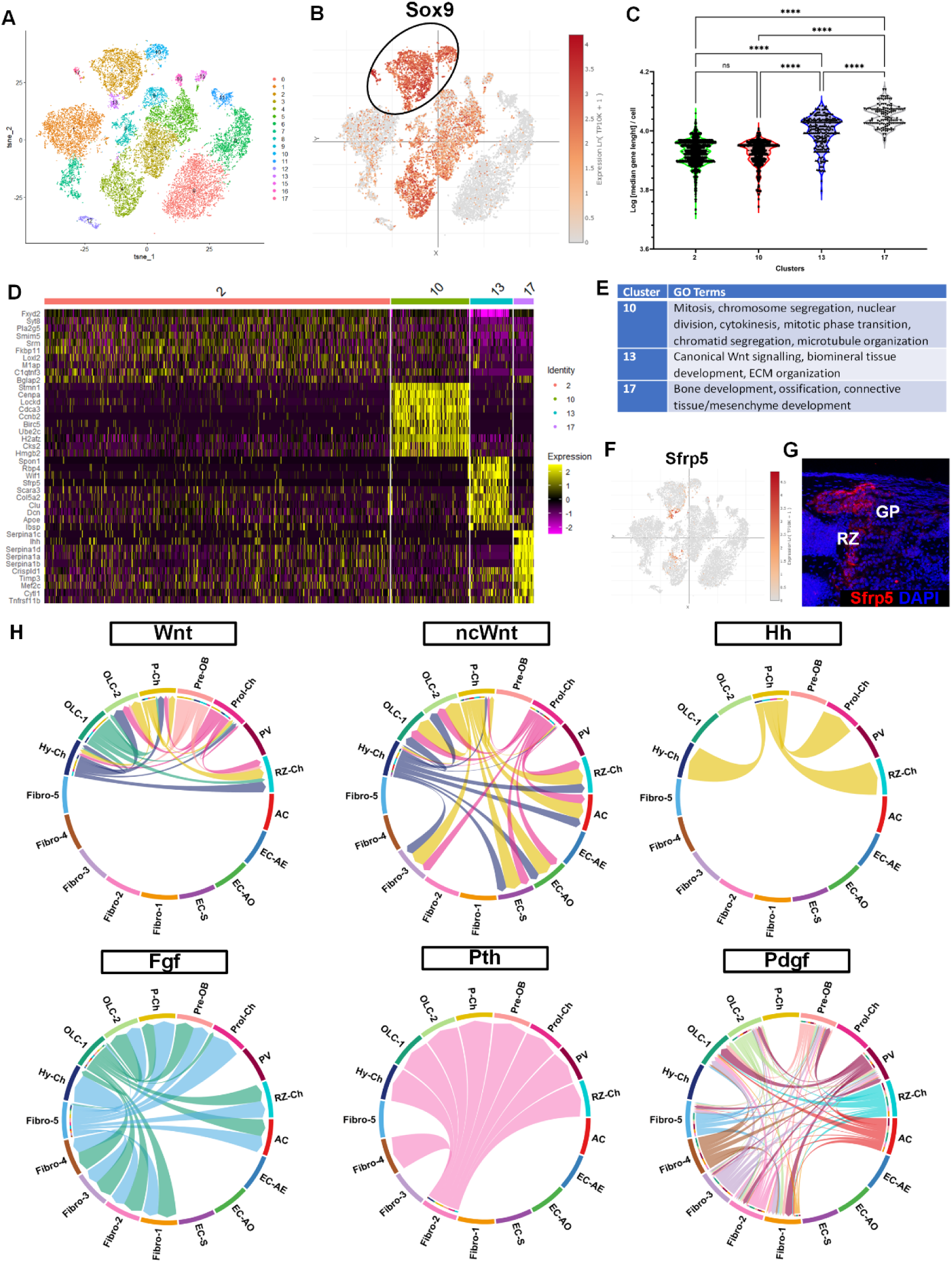
Single cell transcriptomics of mouse postnatal skeletal cells. Open-access **s**ingle cell transcriptomics raw data of postnatal mouse skeletal cells was obtained from previously published data^38^. A) Clustering of scRNA-seq data showing multiple population of skeletal cells. B) Expression of Sox9 transcripts in several clusters of postnatal skeletal cells. The clusters selected for downstream analysis are circled. C) Gene length analysis was perform on transcripts expressed by the four Sox9+ clusters circled in (B). Cells in clusters 13 and 17 express significantly longer transcripts compared to cells from clusters 2 and 10. D) Differential gene expression analysis between the four Sox9+ cell clusters. Heatmap shows the top 10 differentially expressed genes for all clusters. E) GO term analysis was performed on the differentially expressed genes in the four Sox9+ cell clusters. F) Sfrp5, a negative regulator of Wnt signaling, was identified as a differentially expressed gene from cluster 13 with high specificity for this cluster. G) Immunostaining for Sfp5 in P56 mouse femurs confirmed Sfrp5 was specifically expressed in resting zone (RZ) chondrocytes within the distal growth plate (GP). H) CellChat analysis demonstrates molecular crosstalk between several cell types and resting zone chondrocytes, involving mainly Wnt, non-canonical Wnt (ncWnt), Hh, FGF, PTH and PDGF signaling. RZ-Ch: resting zone chondrocytes; AC: articular chondrocytes; EC-AE : endothelial cells – arterial; EC-AO : endothelial cells – arteriolar; EC-S : endothelial cells – sinusoidal; Fibro 1-5: fibroblast-like cells; Hy-Ch: hypertrophic chondrocytes; OLC-1-2: Osteoblast-like cells; P-Ch: perichondral cells; pre-OB: pre-osteoblasts; Prol-Ch: proliferative chondrocytes; PV: perivascular cells.

After clustering the cells using the Louvain algorithm, we performed a pseudotime analysis using the Palantir algorithm, which supported our lineage tracing data and indicated that cells expressing high levels of Sox9 could give rise to cells in the chondrogenic and osteogenic lineages (Fig.6A). Since Sox9+ cells are more plastic during embryonic development (being also capable of forming joint tissues such as ligament, tendons, synovial membrane), we next asked how they differed from postnatal Sox9+ cells. We integrated the postnatal dataset used above with one derived from E16.5 developing limbs^42^. After clustering the integrated data, we found that skeletal cells from both datasets only partially overlapped (Fig.6B). We observed two main clusters of Sox9+ cells that contained both fetal and adult cells (clusters 4 and 5). However, even within those clusters the fetal and adult cells only partially overlapped. We also found several genes that were expressed exclusively in fetal cells, or adult cells (Fig.6C, Supplemental Tables 5-6). This differential gene expression likely explains differences in plasticity between fetal and postnatal Sox9+ cells.

**Figure 6.**
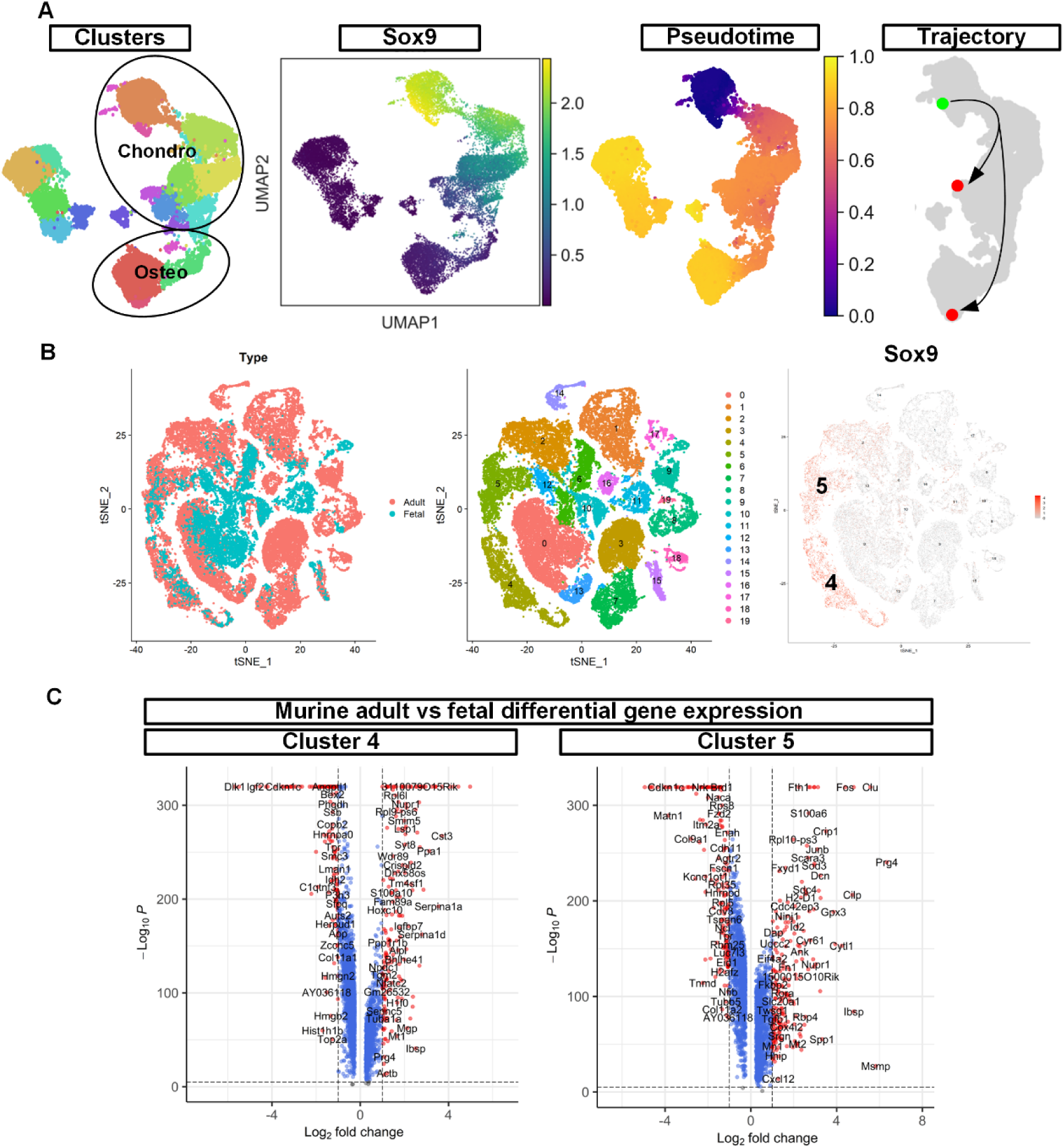
Pseudotime analysis of postnatal mSSCs and comparison with fetal mSSCs. A) We performed graph-based clustering using the Louvain algorithm and pseudotime analysis using the Palantir algorithm. The cellular trajectory computed between Sox9^high^ cells and differentiated osteochondral cells is shown. B) We integrated scRNA-seq data from postnatal (adult) femurs and E16.5 developing mouse limbs. Adult and fetal cells only partially overlapped. Sox9 expression was mainly restricted to clusters 4 and 5 in the integrated dataset. C) In Sox9+ clusters 4 and 5, several genes are upregulated only in fetal or adult Sox9+ cells, reflecting the phenotypical differences observed in Sox9-CreERT lineage tracing experiments between fetal and postnatal mice.

### Sox9+ putative SSCs from postnatal human bones possess multilineage differentiation potential

To determine if postnatal human bones also harbor Sox9+ SSCs, we obtained samples from a pediatric polydactyly patient undergoing amputation of the extra digits. We processed the bones for imaging as for mouse bones. We observed human Sox9 (hSox9) expression in a limited number of cell nuclei in the physeal cartilage surrounding the secondary ossification center. hSox9+ nuclei were enriched in the resting zone of the physis, in subchondral bone, and in articular cartilage (Fig.7A). This was confirmed using a different anti-hSox9 antibody (Fig.S7). hSox9+ cells in subchondral bone and articular cartilage also expressed integrin β5 (ITGB5), which we also observed in some chondrogenic lineage cells in mice (not shown). The fraction of cells positive for hSox9 in the physeal cartilage was measured at 19.5% (Fig.7B).

**Figure 7.**
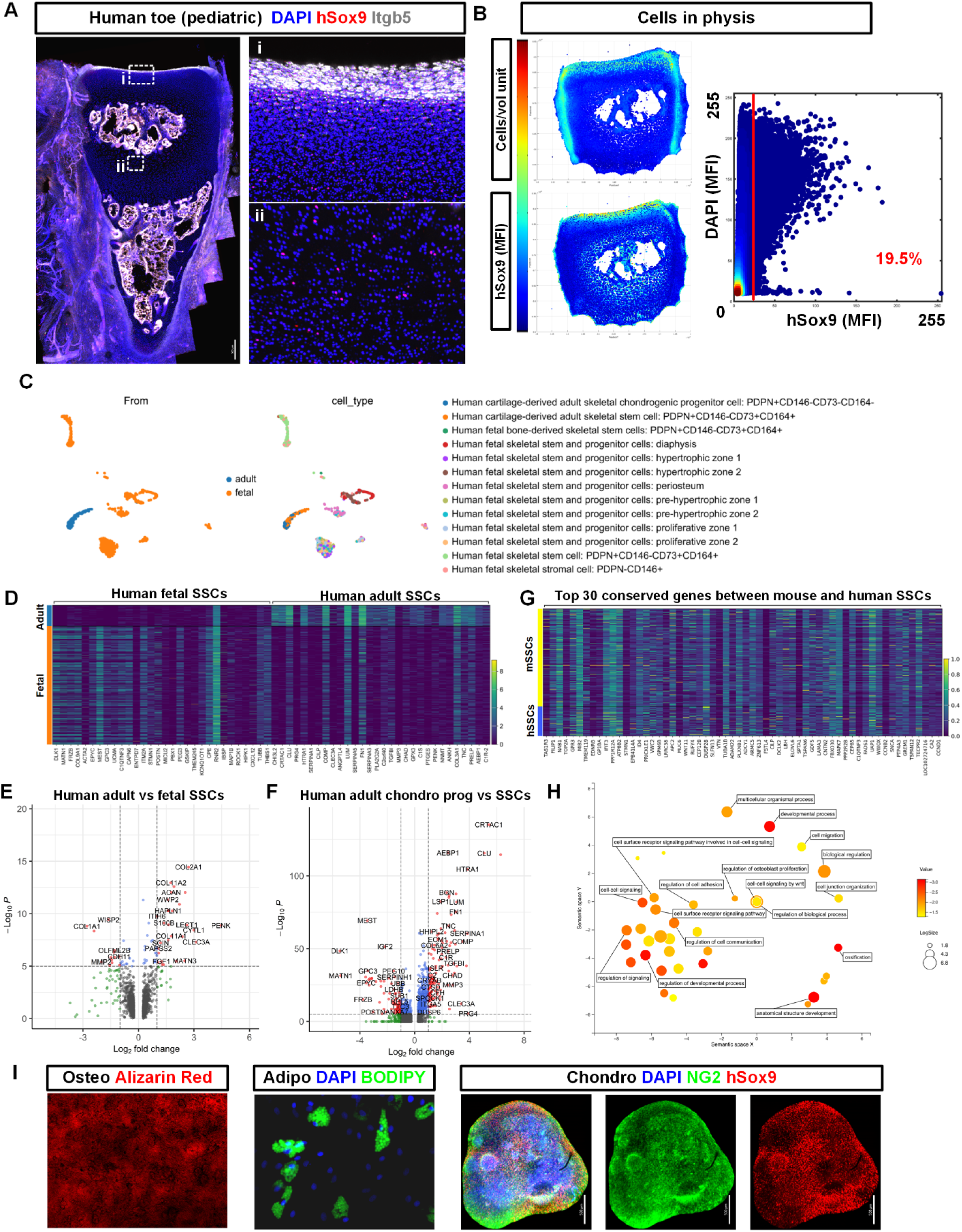
Sox9+ putative osteochondral SSCs in human postnatal skeletal tissue. A) Phalanx of a toe obtained from a pediatric patient with polydactyly whose toe was surgically amputated. The bone was processed similarly to mouse bones, immunostained for hSox9 and ITGB5, and counterstained with DAPI. B) Quantification of hSox9 in cells located in the physis of the bone shown in A. C) Integration of human fetal and adult single cell transcriptomics data indicates that cells derived from adult tissues cluster separately from fetal cells. D) DGE analysis was performed to identify genes that were differentially expressed by adult and fetal hSSCs. E) Volcano plot of the DGE analysis shown in (D) shows that fetal and adult hSSCs share a common gene expression profile that differs only for specific genes. F) DGE analysis of human adult chondrogenic progenitors compared to adult hSSCs used to identify genes involved in self-renewal of hSSCs. G) Gene expression profile comparison between murine and human SSCs used to identify gene involved in self-renewal of both mSSCs and human SSCs. H) GO term analysis of the top 100 expressed genes that are conserved between mSSCs and hSSCs reveals many genes involved in developmental processes, cell signaling, and more specifically Wnt signaling. I) Skeletal cells isolated from patient undergoing epiphysiodesis were expanded for three passages in vitro and subjected to multilineage differentiation assay.

To determine if human SSCs modify their gene expression profile between fetal and postnatal development akin to what was observed in murine cells, we obtained previously published single cell transcriptomics data from human cells at various developmental stages^15^. Similar to what we observe in mice, cells derived from adults clustered separately from fetal cells (Fig.7C). DGE analysis identified several differentially-expressed genes between fetal and adult hSSCs (Fig.7D-E, Supplemental Table 7), including extracellular matrix components as well as growth factors. To identify genes specifically involved in hSSCs self-renewal, we next performed DGE analysis between adult human SSCs and the more committed adult chondrogenic progenitors (Fig.7F, Supplemental Table 8). We found that committed chondrogenic progenitors overexpressed genes such as POSTN, FRZB, DLK1, IGF2 and MEST, whereas hSSCs overexpressed PRG4, MMP3, TGFBI, COL6A2, SERPINA1, FN1, and CLU, amongst others. To further identify genes involved in SSCs self-renewal in both mice and humans, we next compared gene expression between mSSCs and hSSCs. We found that murine and human SSCs share a common gene expression profile (Fig.7G, Fig.S8) with some notable differences (Fig.S9). Specifically, GO term analysis performed on the top 100 conserved genes between mSSCs and hSSCs identified several genes involved in development, cell signaling, and regulation of Wnt signaling (Fig.7H).

Finally, to demonstrate that hSSCs could be isolated while maintaining multipotency in vitro, we next obtained pediatric physis samples from patients undergoing epiphysiodesis for minor limb length discrepancy without underlying genetic causes. Skeletal cells were isolated by enzymatic dissociation and we performed single cell transcriptomics, confirming that ITGB5 could be potentially used as a surrogate marker to purify at least a subpopulation of Sox9+ cells from human postnatal bones (not shown). Skeletal cells from human tissues were then purified by FACS based on ITGB5 expression and culture expanded. After three passages, the human cells were placed in osteogenic or adipogenic media or aggregated and placed in chondrogenic medium. Our results indicate that Sox9+/ITGB5+ cells isolated from postnatal human bones can be purified and expanded ex vivo while maintaining their multilineage differentiation potential (Fig.7I).

## Discussion

Lineage tracing in vivo represents the gold standard to identify populations of putative somatic stem cells during embryonic and postnatal development ^43–45^. Until recently, the identification of bona fide SSCs in postnatal bones has been hindered by technical limitations associated with processing of hard tissues for (immuno-)histological analysis, the autofluorescence of skeletal tissues, and the lack of computational tools to quantify various and abundant subpopulations of cells in entire organs. We recently addressed these hurdles by developing techniques enabling multiplexed 3D fluorescence imaging of large skeletal tissues^33–35^. We also developed computational tools allowing fast and reproducible imaging cytometry of large imaging datasets. Using these techniques, we have demonstrated that the resting zone of the postnatal mouse physis harbors multipotent osteochondral Sox9+ cells that self-renew over the lifespan of the animal driving postnatal bone growth and persist in skeletally mature mice to maintain skeletal tissue homeostasis. These cells are either the progenitors of recently identified putative SSCs in the physeal cartilage or represent a subpopulation of these cells. Our data also builds on and supports previous studies demonstrating that the physis harbors osteochondral stem cells^15,27,31,32,46,47^, that Sox9 labels osteochondral progenitors^26,48^, and that Sox9+ cells in the periosteum participate in injury repair^49,50^. These postnatal Sox9+ SSCs differ from their embryonic or perinatal counterparts in their apparent inability to generate marrow stromal cells expressing Cxcl12^23,26^ or other skeletal tissues in the joints^51–53^. We hypothesize that Sox9+ SSCs in the periphery of the physis are seeded or left behind in the periosteum as the bone elongates, where they remain mostly quiescent until they are activated to proliferate and participate in tissue maintenance, regeneration or repair. To test this hypothesis would require more sophisticated lineage tracing using dual inducible recombinases and/or multiple thymidine analogues incorporation and retention assays. Whether periosteal Sox9+ cells retain their self-renewal potential remains unknown since their abundance fell below detectable limits in long chases.

Our EdU labeling experiments indicate that most periosteal cells are actively dividing in growing, postnatal bones, since many of them incorporated EdU during a short, three-day pulse. However, they likely underwent only one or two rounds of division during that time, after which they stopped proliferating and became postmitotic cortical osteocytes that retained EdU for over 30 days. However, no EdU+ cortical osteocytes were observed in longer chase (90 days), suggesting that these osteocytes had either turnover and been replaced by non-labeled cells, or had resumed proliferation and lost the EdU label, which is improbable. On the other hand, tdT+ cells derived from Sox9+ SSCs were much rarer in periosteum and cortical bone but their number increased over time, replacing the cells that had incorporated EdU. This strongly suggests that Sox9+ cells are the upstream progenitors of the proliferative periosteal cells and that cortical bone tissue in growing mouse bones exhibits a high rate of tissue turnover.

Our single cell transcriptomics analyses revealed the importance of the regulation of Wnt signaling in mSSCs, since these cells express many negative regulators of Wnt including Wif1 and Sfrp5. While the role of Wnt signaling in skeletal development has long been described, recent studies also indicated a role for Wnt signaling in SSCs ^15,27,40^. More specifically, Wnt signaling appears to promote chondrocyte proliferation and column formation for longitudinal bone growth using the planar cell polarity pathway^40^. Our results support this conclusion that SSCs suppress Wnt activity to maintain their quiescent state in the resting zone. How other signaling pathways such as Hh, FGF, and PTH regulate SSCs self-renewal remains unclear. Our analyses on human SSCs indicated that Wnt signaling was also likely a key molecular player in their fate decisions. The small differences observed between murine and human SSCs can be explained by differences in the postnatal developmental stages between the samples.

Future research confirming the persistence of Sox9+ SSCs in mature adult humans is required; however, our preliminary results with human growth plate tissue suggests that this may be the case. Whether such populations would be present in long bones in which the growth plate closes (i.e. in humans, but not rodents where the growth plate remains open) is another question that will need to be answered. This will be particularly important, given that the majority of chronic musculoskeletal disease affects the aging adult population.

To summarize, our study provides evidence of a novel population of self-renewing osteochondral SSCs in the postnatal murine and human skeleton. These findings are key to our understanding of skeletal stem cells, and a necessary step towards the future development of innovative stem cell-based regenerative therapies for various orthopedic conditions.

### Limitations of the study

Our lineage tracing of Sox9-CreERT expressing cells combined with analysis of Sox9 protein expression and EdU incorporation/retention provides the first tangible experimental evidence of in vivo self-renewal in any type of SSCs identified to date. However, our findings on self-renewal and multipotency are limited to the population level. While our data confirms that a subpopulation of Sox9+ cells possess self-renewal capacity, it remains possible that these Sox9+ SSCs in the resting zone could be further divided in several subpopulations of unipotent cells. Since we observed tdT labeling of osteoblasts in the metaphysis, we cannot exclude the possibility that these contained self-renewing osteoblast-restricted cells. and that resting zone Sox9+ cells only give rise to chondrocytes. However, such a population of self-renewing osteoblast-like cells has never been described, we showed that lineage tracing of cell downstream of Sox9+ cells does not show any indication of self-renewal in osteoblasts (Fig.S10), and our EdU incorporation/retention data (Fig.1-2) does not show evidence of self-renewal in osteoblast-lineage cells, which proliferated fast and turned-over rapidly. Moreover, it is widely assumed that up to 60% of postnatal osteoblasts originate from growth plate chondrocytes^54–56^ In our lineage tracing experiments, we observed some discrepancies between the expression of tdT and the Sox9 protein two-days post-labeling. Indeed, several osteogenic tdT+ cells had no detectable Sox9 protein whereas some Sox9+ proliferative chondrocytes did not get labeled with tdT. It is possible that the short two-day chase was enough for some Sox9+ cells to commit to the osteoblast lineage and downregulate Sox9. Similarly, it is possible that proliferative chondrocytes stopped transcribing the Sox9 gene but still contained detectable Sox9 protein. An alternative explanation would involve a combination of differential transcriptional bursting pattern, protein maturation, and protein stability between the endogenous Sox9 gene, the Sox9-CreERT transgene, and the Rosa26 tdTomato Cre-reporter and their respective encoded proteins. It is also possible that given the short tamoxifen pulses used and the short bioavailability of its active metabolites, cells located deep within the avascular cartilage were not exposed to sufficient levels of tamoxifen. Regardless, none of these limitations and hypotheses alter our lineage tracing interpretation.

Another intriguing finding resides in the population kinetics of tdT+ and tdT+Sox9+ cells we observed. In growing bones, we observed a biphasic population kinetics of tdT+ cells, where their number increased within the first 30 days and then decreased back to their initial numbers in longer chases. The number of tdT+Sox9+ cells (putative self-renewing SSCs) did not follow the same kinetics. In addition, the population kinetics of these cell types was also different in mature bones. The number of transiently amplifying clones that expressed Sox9 (e.g. proliferative chondrocytes, pre-osteoblasts, etc.) in growing bones combined with the increasing tissue volume and mass over time makes the interpretation of this data challenging. Moreover, it is unclear what fraction of the total Sox9+ SSCs we labeled using such short labeling pulses, further adding the challenge of interpreting this type of data. To fully reconcile these findings will require additional studies and computational modelling approaches.

Our data clearly shows that Sox9+ SSCs are maintained in skeletally mature animals and continue to contribute to bone homeostasis. Isolating SSCs from older animals proved challenging and it is possible that their growth requirements changed, that older SSCs are more prone to cellular senescence, or that we observed age-related stem cell exhaustion or loss of function. Further studies will be required to address these issues.

## Acknowledgments

The authors would like to thank the Research Chair in Regenerative Orthopedic surgery (The Ottawa Hospital) and Stem Cell Network for funding to DLC, the Hans K. Uhthoff graduate scholarship for funding to SF and BT, and the Canadian Institutes of Health Research for funding to SS. The authors acknowledge the Cell Biology and Image Acquisition Core (RRID: SCR_021845) funded by the University of Ottawa, Natural Sciences and Engineering Research Council of Canada, and the Canada Foundation for Innovation, and thank them for their support.

## Methods

### Animals

All experiments in mice were performed in accordance with the Animal Care and Veterinary Service (ACVS) guidelines at University of Ottawa and approved by the institutional ethics review board. The Sox9-CreERT;Ai14 double mutant mouse line used in this study was obtained by breeding two mouse strains, both in the C57Bl/6 background: 1) Sox9-cre/ERT2)1Msan/J; MGI: 5009223), and 2) Rosa26 LoxP-STOP-LoxP fluorescent Cre-reporter mouse line in which expression of the tdTomato (Ai14) reporter is induced by Cre activation (B6.Cg-Gt(ROSA)26Sortm14(CAG-tdTomato)Hze/J; MGI : J:155793)2. Both strains were purchased from the Jackson Laboratory. Experiments were conducted using both males and females whenever possible. Two age groups of male and female mice were used for this study, juvenile mice (P56) and adult mice (P180). Once they reached their appropriate age, they were injected 1-3 times with 75mg/kg body weight 4-hydroxytamoxifen (4OHT) or tamoxifen (TAM), as indicated in the text/figures, at 24h intervals intraperitoneally (i.p) to activate CreERT and induce the expression of tdTomato in Sox9+ cells. At designated time-points following 4OHT/TAM administration (TAM at P56 or P180, timepoints P58 or P182, P86 or P210, and P236 or P360), the mice were euthanized by CO2 inhalation followed by cervical dislocation.

### Skeletal cells isolation

Sox9-CreERT;Ai14 mice were injected at P56 or P180 (as indicated in text/ figures) with a single dose of 4OHT, then femurs or joints were harvested after 48h. Epiphyses were cut, marrow was flushed out using 25G needle filled with PBS, then mechanically dissociated by passing through 25G needle. Bone was cut in small pieces, enzymatically dissociated, then crushed using a sterile mortar and pestle under the cell culture hood. Stemxyme (Worthington Biochemical Corp.) was used for enzymatic digestion. Alternatively, a custom enzymatic cocktail (as indicated in text/ figure) was used containing PBS with 2% FBS, 2.5 mg/mL of collagenase I (Worthington), 0.7 mg/mL of collagenase II (Worthington), 1 U/mL of dispase (Worthington) and 5uM of Ca2+(Sigma-Aldrich). The samples were then placed on a shaker for 45 min at 37°C. Cells were filtered using a 100um cell strainer, then plated in cell culture dishes with murine Mesencult expansion medium (StemCell Technologies). Skeletal cells were passaged biweekly or when they reached 80% confluency, whichever came first. For enrichment, negative selection was used where CD31+ endothelial, Ter119+ erythroid, and CD45+ hematopoietic cells were magnetically depleted using the EasySep™ Mouse Mesenchymal Stem/Progenitor Cell Enrichment Kit StemCell Technologies (customized by adding a biotinylated CD31 antibody).

### Antibodies

The list of primary antibodies used in this study are summarized in supplemental Table 1. AlexaFluor-conjugated secondary antibodies were obtained from Thermo. All primary and secondary antibodies were reconstituted according to manufacturer’s instructions (when required), then diluted 1:1 in glycerol and stored at -20°C.

### Colony forming assay

Sox9-CreERT;Ai14 mice were injected at P56 or P180 (as indicated in text/ figures) with a single dose of 4OHT. Femurs or joints were harvested two days post-4OHT, and cells were isolated as described in section 2.3. Cells were then counted and seeded at densities of 1.2 × 10^5^ cells/cm^2^, in triplicates, in 24 well-plates, in mouse Mesencult Expansion medium (StemCell Technologies). After 10 days in culture, cells were fixed and stained with 0.5% crystal violet in methanol. The total number of colonies and the number of tdT+ colonies were counted (colonies containing >20 fibroblastoid cells were counted) using an Epifluorescence microscope (Nikon eclipse TE2000-E, filter B-2E/C , DM:505, Excitation: 465-495 and Emission: 515-555, CFI Plan Fluor 10X Dry N.A 1.3).

### In vitro differentiation

Sox9-CreERT;Ai14 mice were injected at P56 or P180 (as indicated in text/figures) with a single dose of 4OHT, then femurs or joints were harvested two days post-4OHT, and cells were isolated as described in section 2.3, then cultured in Mesencult medium (StemCell Technologies). Cells were expanded until confluence in 9 wells (three biological replicates with three technical replicates each) of 24-well plates. Adipogenic, chondrogenic and osteogenic differentiation was induced in three wells each. For chondrogenic differentiation, cells were pelleted by centrifugation at 1200RPM using a centrifuge (Eppendorf, 5810 R, v8.4) in 15 mL falcon tubes to promote aggregation. Serum-free chondrogenic medium contained 10ng/mL TGF-β1, 100nM dexamethasone, 50 µg/mL ascorbic acid-2-phosphate, 100 µg/mL sodium pyruvate, 40 µg/mL L-proline, 1X ITS+3 (Sigma) and 1.25mg/mL Bovine Serum Albumin. For osteogenic and adipogenic differentiation, cells were differentiated using mouse MesenCult™ Adipogenic and Osteogenic Differentiation Kits from StemCell Technologies. Cells were differentiated for 3 weeks and then fixed, immunostained, as described in section 2.6, for differentiation markers and imaged using confocal microscopy. Alternatively, mineralized matrix was stained with 2% m/v alizarin Red diluted in ddH2O, and adipocytes with 2uM BODIPY™ 493/503 (4,4-Difluoro-1,3,5,7,8-Pentamethyl-4-Bora-3a,4a-Diaza-s-Indacene) (Invitrogen).

### Flow cytometry

To sort tdT+ cells, Sox9-CreERT;Ai14 mice were injected at P56 with a single dose of 4OHT, then joints were harvested two days post-4OHT. Joints were digested and depleted from hematopoietic and endothelial cells as described above. tdT+ cells were isolated from both distal femurs and proximal tibias using a MoFlo XDP flow cytometer and sorter located at the Flow cytometry and Cell sorting facility of the Ottawa Hospital Research Institute

### Single cell transcriptomics

The postnatal single cell transcriptomics data used was previously published^38^ and downloaded from https://singlecell.broadinstitute.org/single_cell/study/SCP361/mouse-bone-marrow-stroma-in-homeostasis#study-summary. The data underwent preprocessing steps, including scaling, log normalization, and processing with Seurat. For cluster annotation, identities were assigned based on the original clustering. Additionally, a CellChat^41^ analysis was conducted on the initial data to identify the major pathways regulating Sox9 high clusters (cluster 2, 10, 13, and 17). Subsequently, Sox9 high clusters were subsetted, and differential gene expression (DGE) analysis was performed using the Wilcoxon test to identify the top markers for each cluster, which were visualized in a heatmap. To understand the functional significance of these top genes, a Gene Ontology (GO) enrichment analysis was executed, focusing on genes expressed in at least 50% of the cluster.

For comparing adult and developing mouse skeletal cells, raw publicly available data from adult (GSM3674239_b1, GSM3674240_b2, GSM3674241_b3, GSM3674242_b4) and E16.5 mouse (GSM4748807)^42^ samples were merged, filtered, scaled, and log normalized using Seurat. The data dimensionality was reduced to the top 50 principal components (PCA), followed by graph-based clustering using the Louvain algorithm. To mitigate batch effects, data integration was performed using the Seurat Integration function, and subsequent steps included scaling, log normalization, reducing to the top 50 PCA components, and clustering based on the top 20 components for batch correction control. Visualizations using UMAP and t-SNE were generated. Sox9 high clusters (cluster 2, 4, and 5) were selected for further analysis, where conserved markers and differentially expressed markers were identified and visualized using ggplot.

For pseudotime analysis, the data was log normalized and processed using Seurat. The data was reduced to the top 50 PCA components and graph-based clustering was performed using the Louvain algorithm^57^. The clusters were visualized on 2D map computed by UMAP. Pseudotime analysis was performed using the Palantir algorithm^58^.

For the Smart-seq2 human dataset, the raw unmapped data were downloaded from GEO SRA PRJNA478935 and processed as previously described^59^. However, reads were aligned to the human reference genome GRCh38 using STAR v2.7.11a with default parameters^60^. Gene expression was calculated using RSEM v1.3.3^61^, and only the transcripts per million (TPM) matrix output was used for downstream analysis in both Seurat v4.4.0^62^ and Scanpy v1.9.5^63^, as described in Seurat’s vignette^64^ and Scanpy’s vignette^65^. In both Seurat and Scanpy, default parameters for quality control (QC) analysis of single-cell RNA-sequencing (scRNA-seq) data were applied. Cells with fewer than 200 genes, more than 2,500 genes, or mitochondrial gene content higher than 10% were removed. Additionally, genes identified in fewer than 3 cells were also removed. The analysis then proceeded with typical normalization, dimensionality reduction (PCA and UMAP), and cell clustering. Subsequently, MAST^66^ was utilized for differential gene expression analysis and marker gene identification. Cell types were annotated based on each dataset’s original authors.

For the human–mouse cross-species data integration, both SATURN^67^ and TACTiCS^68^ were applied independently using their default parameters. Batch effects were assessed and corrected using scVI v1.0.3^69^. Finally, lvm-DE^70^ was utilized for differential gene expression analysis.

### Tissue harvesting and processing for immunofluorescence

Bones were processed as described^33,35^. Briefly, Sox9-CreERT;Ai14 mice were injected at P56 or P180 with one or three injections of tamoxifen or 4-OHT (as indicated in text/figures), and femurs or joints were collected at specific timepoints ( two, 30 and 180 days post-4OHT injection), for both conditions (healthy versus FOD injury) and both gender ( three males and three females/condition). Femurs and joints were cleaned from muscle tissue then fixed overnight in 16% methanol-free formaldehyde (Electron Microscopy Sciences) diluted to 4% in PBS at 4°C with rotation. This was followed by decalcification for two weeks in 10% EDTA (Sigma) diluted in PBS, pH 8, at 4°C with stirring.

### Vibratome sectioning

Decalcified bones/joints were embedded in 4% low-temperature gelling agarose (high gel strength grade, Bioshop, CAS#9012-36-6), and then cut into 300-500 µm-thick sections using a Leica VT1200S vibratome with low profile, injector-type Endurium zirconia ceramic blades (Cadence Inc.). These sections were then probed by immunofluorescence staining and processed for confocal microscopy.

### Immunofluorescence

Femur/joint sections were blocked and permeabilized for 1h with blocking buffer made from TBS (final concentration 0.1M Tris, 0.15 M NaCl, pH 7.5), 0.05% Tween-20, 20% DMSO (all from Sigma), 5% donkey serum (Jackson ImmunoResearch) and 0.3% Triton X-100 (Sigma). Sections were then stained with primary antibodies (refer to section 2.13 table x for working dilutions) in blocking buffer overnight. Sections were washed 5×60 minutes with TBS containing 0.05% Tween-20, stained with fluorophore-conjugated secondary antibodies (1:200 dilution) and counterstained with DAPI (1:500 dilution of a 2mg/mL stock solution), in blocking buffer, overnight. The fluorophores used were AlexaFluor 555, 488, 633, 680, with or without biotin-streptavidin amplification, and nuclei were counterstained with DAPI (Thermo). The samples were then washed 5×60 minutes as above, then optically cleared (see below). All steps were done at room temperature with gentle shaking.

### EdU proliferation assay

Sox9-CreERT;Ai14 mice were injected with one or three injections of 4OHT, and simultaneously with one, three or five EdU injections (2.5mg/mL) at 24h interval (as indicated in text/figures). Femurs were harvested and processed as described above. Femur sections were blocked then stained with Click-iT EdU Alexa Fluor 488 imaging kit (Invitrogen) overnight. The next day, sections were stained as described above.

### Optical clearing and mounting of sections

For optical clearing, we used a modification of the Ce3D protocol^71^. This method is faster than our previously published TDE-based clearing method^33,35^ and comparable in terms of clearing efficiency (Fig.S11). Briefly, femur samples were optically cleared with histodenz (Sigma). Stained sections were incubated in 88% histodenz overnight at room temperature with gentle shaking. The final mounting medium consisted of 88% histodenz dissolved in TBS with 0.1% tween-20, 0.01% NaN3, pH=8.5. The refractive index was measured using a handheld refractometer (Atago) and adjusted to 1.467 with histodenz. Sections were then mounted on Superfrost microscope slides using custom-designed silicone spacers (Grace Biolabs) and size 1.5 coverslips.

### Confocal microscopy

Imaging was performed on a confocal microscope (Leica TCS SP8) equipped with three PMTs (photomultiplier tubes), two HyD detectors and five laser lines: blue diode (405 nm), argon (458, 476, 488, 496 and 514 nm) and three helium neon (543, 594 and 633 nm). Type G immersion liquid (Leica) was used with an RI of 1.467. Image acquisition was done using a 20× multiple immersion lens (NA 0.75, FWD 0.680 mm). All scans were acquired at room temperature with a speed of 600 Hz, in the bidirectional mode and with z-spacing of 2.5 um. Images were acquired in 8-bit format with a 1.0× zoom at 1,024 × 1,024 resolution.

### Analysis and statistics

Images were compressed (lossless), visualized, and segmented using Imaris v9.8.2 (Bitplane) and segmentation data were exported for downstream analyses using XiT software. For statistical analyses, gated data was exported from XiT and analyzed using GraphPad Prism v9. For statistical analyses, data from all technical and biological replicates was compiled for each age, sex group and condition, and plotted (mean +/- SD), unless otherwise specified. For each cohort, an outlier analysis was performed, and normality tested using the Shapiro-Wilks test. Each age, sex group and condition were compared using two-way ANOVA with significance threshold set at p=0.05. Where relevant, the n and p values are indicated in the text or figure legends.

## References

1. Coutu, D. L. & Galipeau, J. Molecular and endocrine mechanisms underlying the stem cell theory of aging. in Adult Stem Cells (ed. Turksen, K.) 389–417 (Springer, 2014). doi:10.1007/978-1-4614-9569-7.

2. Ermolaeva, M., Neri, F., Ori, A. & Rudolph, K. L. Cellular and epigenetic drivers of stem cell ageing. Nat Rev Mol Cell Biol 19, 594–610 (2018).

3. Ambrosi, T. H. et al. Aged skeletal stem cells generate an inflammatory degenerative niche. Nature (2021) doi:10.1038/s41586-021-03795-7.

4. Coutu, D. L., François, M. & Galipeau, J. Inhibition of cellular senescence by developmentally regulated FGF receptors in mesenchymal stem cells. Blood 117, 6801–6812 (2011).

5. Freund, A., Orjalo, A. V., Desprez, P. Y. & Campisi, J. Inflammatory networks during cellular senescence: causes and consequences. Trends Mol Med 16, 238–246 (2010).

6. Despars, G., Carbonneau, C. L., Bardeau, P., Coutu, D. L. & Beauséjour, C. M. Loss of the Osteogenic Differentiation Potential during Senescence Is Limited to Bone Progenitor Cells and Is Dependent on p53. PLoS One 8, 1–11 (2013).

7. Park, D. et al. Endogenous bone marrow MSCs are dynamic, fate-restricted participants in bone maintenance and regeneration. Cell Stem Cell 10, 259–272 (2012).

8. Morikawa, S. et al. Prospective identification, isolation, and systemic transplantation of multipotent mesenchymal stem cells in murine bone marrow. J Exp Med 206, 2483–2496 (2009).

9. Bonyadi, M. et al. Mesenchymal progenitor self-renewal deficiency leads to age-dependent osteoporosis in Sca-1/Ly-6A null mice. Proceedings of the National Academy of Sciences 100, 5840–5845 (2003).

10. Suire, C., Brouard, N., Hirschi, K. & Simmons, P. J. Isolation of the stromal-vascular fraction of mouse bone marrow markedly enhances the yield of clonogenic stromal progenitors. Blood 119, (2012).

11. Sacchetti, B. et al. Self-Renewing Osteoprogenitors in Bone Marrow Sinusoids Can Organize a Hematopoietic Microenvironment. Cell 131, 324–336 (2007).

12. Coutu, D. L., François, M. & Galipeau, J. Inhibition of cellular senescence by developmentally regulated FGF receptors in mesenchymal stem cells. Blood 117, (2011).

13. Kunisaki, Y. et al. Arteriolar niches maintain haematopoietic stem cell quiescence. Nature 502, 637–643 (2013).

14. Pinho, S. et al. PDGFRα and CD51 mark human Nestin ^+^ sphere-forming mesenchymal stem cells capable of hematopoietic progenitor cell expansion. J Exp Med 210, 1351–1367 (2013).

15. Chan, C. K. F. et al. Identification of the Human Skeletal Stem Cell. Cell 175, 43–56.e21 (2018).

16. Chan, C. K. F. et al. Endochondral ossification is required for haematopoietic stem-cell niche formation. Nature 457, 490–494 (2009).

17. Méndez-Ferrer, S. et al. Mesenchymal and haematopoietic stem cells form a unique bone marrow niche. Nature 466, 829–834 (2010).

18. Greenbaum, A. et al. CXCL12 in early mesenchymal progenitors is required for haematopoietic stem-cell maintenance. Nature 495, 227–230 (2013).

19. Qian, H. et al. Molecular Characterization of Prospectively Isolated Multipotent Mesenchymal Progenitors Provides New Insight into the Cellular Identity of Mesenchymal Stem Cells in Mouse Bone Marrow. Mol Cell Biol 33, 661–677 (2012).

20. Maes, C. et al. Osteoblast precursors, but not mature osteoblasts, move into developing and fractured bones along with invading blood vessels. Dev Cell 19, 329–344 (2010).

21. Chen, J. et al. Osx-Cre targets multiple cell types besides osteoblast lineage in postnatal mice. PLoS One 9, 1–6 (2014).

22. Liu, Y. et al. Osterix-Cre Labeled Progenitor Cells Contribute to the Formation and Maintenance of the Bone Marrow Stroma. PLoS One 8, (2013).

23. Mizoguchi, T. et al. Osterix marks distinct waves of primitive and definitive stromal progenitors during bone marrow development. Dev Cell 29, 340–349 (2014).

24. Zhou, B. O., Yue, R., Murphy, M. M., Peyer, J. & Morrison, S. J. Leptin Receptor-expressing mesenchymal stromal cells represent the main source of bone formed by adult bone marrow. Cell Stem Cell 15, 156–168 (2014).

25. Ding, L., Saunders, T. L., Enikolopov, G. & Morrison, S. J. Endothelial and perivascular cells maintain haematopoietic stem cells. Nature 481, 457–462 (2012).

26. Ono, N., Ono, W., Nagasawa, T. & Kronenberg, H. M. A subset of chondrogenic cells provides early mesenchymal progenitors in growing bones. Nat Cell Biol 16, 1157–1167 (2014).

27. Newton, P. T. et al. A radical switch in clonality reveals a stem cell niche in the epiphyseal growth plate. Nature 567, 234–238 (2019).

28. Worthley, D. L. et al. Gremlin 1 identifies a skeletal stem cell with bone, cartilage, and reticular stromal potential. Cell 160, 269–284 (2015).

29. Mizuhashi, K. et al. Resting zone of the growth plate houses a unique class of skeletal stem cells. Nature 563, 254–258 (2018).

30. Debnath, S. et al. Discovery of a periosteal stem cell mediating intramembranous bone formation. Nature 562, 133–9 (2018).

31. Chan, C. K. F. et al. Identification and specification of the mouse skeletal stem cell. Cell 160, 285– 298 (2015).

32. Chagin, A. S. & Newton, P. T. Postnatal skeletal growth is driven by the epiphyseal stem cell niche: potential implications to pediatrics. Pediatric Research vol. 87 986–990 Preprint at 10.1038/s41390-019-0722-z (2020).

33. Kunz, L. & Coutu, D. L. Multicolor 3D Confocal Imaging of Thick Tissue Sections. Methods in Molecular Biology 2350, 95–104 (2021).

34. Coutu, D. L., Kokkaliaris, K. D., Kunz, L. & Schroeder, T. Three-dimensional map of nonhematopoietic bone and bone-marrow cells and molecules. Nat Biotechnol 35, (2017).

35. Coutu, D. L., Kokkaliaris, K. D., Kunz, L. & Schroeder, T. Multicolor quantitative confocal imaging cytometry. Nat Methods 15, (2018).

36. Karsenty, G., Kronenberg, H. M. & Settembre, C. Genetic Control of Bone Formation. (2009) doi:10.1146/annurev.cellbio.042308.113308.

37. Felker, A. et al. In vivo performance and properties of Tamoxifen metabolites for CreERT2 control. PLoS One 11, 1–17 (2016).

38. Baryawno, N. et al. Resource A Cellular Taxonomy of the Bone Marrow Stroma in Resource A Cellular Taxonomy of the Bone Marrow Stroma in Homeostasis and Leukemia. Cell 1–18 (2019) doi:10.1016/j.cell.2019.04.040.

39. Chakra, M. A., Isserlin, R., Tran, T. N. & Bader, G. D. Control of tissue development and cell diversity by cell cycle-dependent transcriptional filtering. Elife 10, (2021).

40. Hallett, S. A. et al. Chondrocytes in the resting zone of the growth plate are maintained in a wnt-inhibitory environment. Elife 10, (2021).

41. Jin, S. et al. Inference and analysis of cell-cell communication using CellChat. Nature Communications 2021 12:1 **12**, 1–20 (2021).

42. Arostegui, M., Wilder Scott, R., Böse, K. & Michael Underhill, T. Cellular taxonomy of Hic1+ mesenchymal progenitor derivatives in the limb: from embryo to adult. Nature Communications 2022 13:1 **13**, 1–20 (2022).

43. Fox, D. T., Morris, L. X., Nystul, T. & Spradling, A. C. Lineage analysis of stem cells. in StemBook [Internet] 1–18 (2009). doi:10.3824/stembook.1.33.1.

44. Kretzschmar, K. & Watt, F. M. Lineage tracing. Cell 148, 33–45 (2012).

45. Blanpain, C. & Simons, B. D. Unravelling stem cell dynamics by lineage tracing. Nat Rev Mol Cell Biol 14, 489–502 (2013).

46. Shi, Y. et al. Gli1+ progenitors mediate bone anabolic function of teriparatide via Hh and Igf signaling. Cell Rep 36, 109542 (2021).

47. Mizuhashi, K. et al. Resting zone of the growth plate houses a unique class of skeletal stem cells. Nature (2018) doi:10.1038/s41586-018-0662-5.

48. Balani, D. H., Ono, N. & Kronenberg, H. M. Parathyroid hormone regulates fates of murine osteoblast precursors in vivo. Journal of Clinical Investigation 127, 3327–3338 (2017).

49. He, X. et al. Sox9 positive periosteal cells in fracture repair of the adult mammalian long bone. Bone 103, 12–19 (2017).

50. Kuwahara, S. T. et al. Sox9+ messenger cells orchestrate large-scale skeletal regeneration in the mammalian rib. Elife 8, 1–21 (2019).

51. Soeda, T. et al. Sox9-expressing precursors are the cellular origin of the cruciate ligament of the knee joint and the limb tendons. Genesis 48, 635–644 (2010).

52. Blitz, E., Sharir, A., Akiyama, H. & Zelzer, E. Tendon-bone attachment unit is formed modularly by a distinct pool of Scx-and Sox9-positive progenitors. Development (Cambridge) 140, 2680–2690 (2013).

53. Sugimoto, Y. et al. Scx+/Scx9+ progenitors contribute to the establishment of the junction between cartilage and tendon/ligament. Development (Cambridge) 140, 2280–2288 (2012).

54. Park, J. et al. Dual pathways to endochondral osteoblasts: A novel chondrocytederived osteoprogenitor cell identified in hypertrophic cartilage. Biol Open 4, 608–621 (2015).

55. Yang, L., Tsang, K. Y., Tang, H. C., Chan, D. & Cheah, K. S. E. Hypertrophic chondrocytes can become osteoblasts and osteocytes in endochondral bone formation. Proc Natl Acad Sci U S A 111, 12097–12102 (2014).

56. Zhou, X. et al. Chondrocytes Transdifferentiate into Osteoblasts in Endochondral Bone during Development, Postnatal Growth and Fracture Healing in Mice. PLoS Genet 10, (2014).

57. Blondel, D., Guillaume, J., Lambiotte, R. & Lefebvre, E. Fast unfolding of communities in large networks. J. Stat. Mech. P10008 (2008).

58. Setty, M. et al. Characterization of cell fate probabilities in single-cell data with Palantir. Nat Biotechnol 37, 451–460 (2019).

59. Smart-seq2 Single Sample Overview | WARP. https://broadinstitute.github.io/warp/docs/Pipelines/Smart-seq2_Single_Sample_Pipeline/README/.

60. Dobin, A. et al. STAR: ultrafast universal RNA-seq aligner. Bioinformatics 29, 15–21 (2013).

61. Li, B. & Dewey, C. N. RSEM: Accurate transcript quantification from RNA-Seq data with or without a reference genome. BMC Bioinformatics 12, 1–16 (2011).

62. Hao, Y. et al. Integrated analysis of multimodal single-cell data. Cell 184, 3573–3587.e29 (2021).

63. Wolf, F. A., Angerer, P. & Theis, F. J. SCANPY: Large-scale single-cell gene expression data analysis. Genome Biol 19, 1–5 (2018).

64. Seurat - Guided Clustering Tutorial • Seurat. https://satijalab.org/seurat/articles/pbmc3k_tutorial.

65. Preprocessing and clustering 3k PBMCs — scanpy-tutorials 1.4.7.dev38+g3e4e434 documentation. https://scanpy-tutorials.readthedocs.io/en/latest/pbmc3k.html.

66. Finak, G. et al. MAST: A flexible statistical framework for assessing transcriptional changes and characterizing heterogeneity in single-cell RNA sequencing data. Genome Biol 16, 1–13 (2015).

67. Rosen, Y. et al. Towards Universal Cell Embeddings: Integrating Single-cell RNA-seq Datasets across Species with SATURN. bioRxiv 2023.02.03.526939 (2023) doi:10.1101/2023.02.03.526939.

68. Biharie, K., Michielsen, L., Reinders, M. J. T. & Mahfouz, A. Cell type matching across species using protein embeddings and transfer learning. Bioinformatics 39, i404–i412 (2023).

69. Gayoso, A. et al. A Python library for probabilistic analysis of single-cell omics data. Nature Biotechnology 2022 40:2 **40**, 163–166 (2022).

70. Boyeau, P. et al. An empirical Bayes method for differential expression analysis of single cells with deep generative models. Proc Natl Acad Sci U S A 120, e2209124120 (2023).

71. Li, W., Germain, R. N. & Gerner, M. Y. High-dimensional cell-level analysis of tissues with Ce3D multiplex volume imaging. Nat Protoc 14, 1708–1733 (2019).

