## Supplementary material for "Self-renewing Sox9+ osteochondral stem cells in the postnatal skeleton": Fig.S

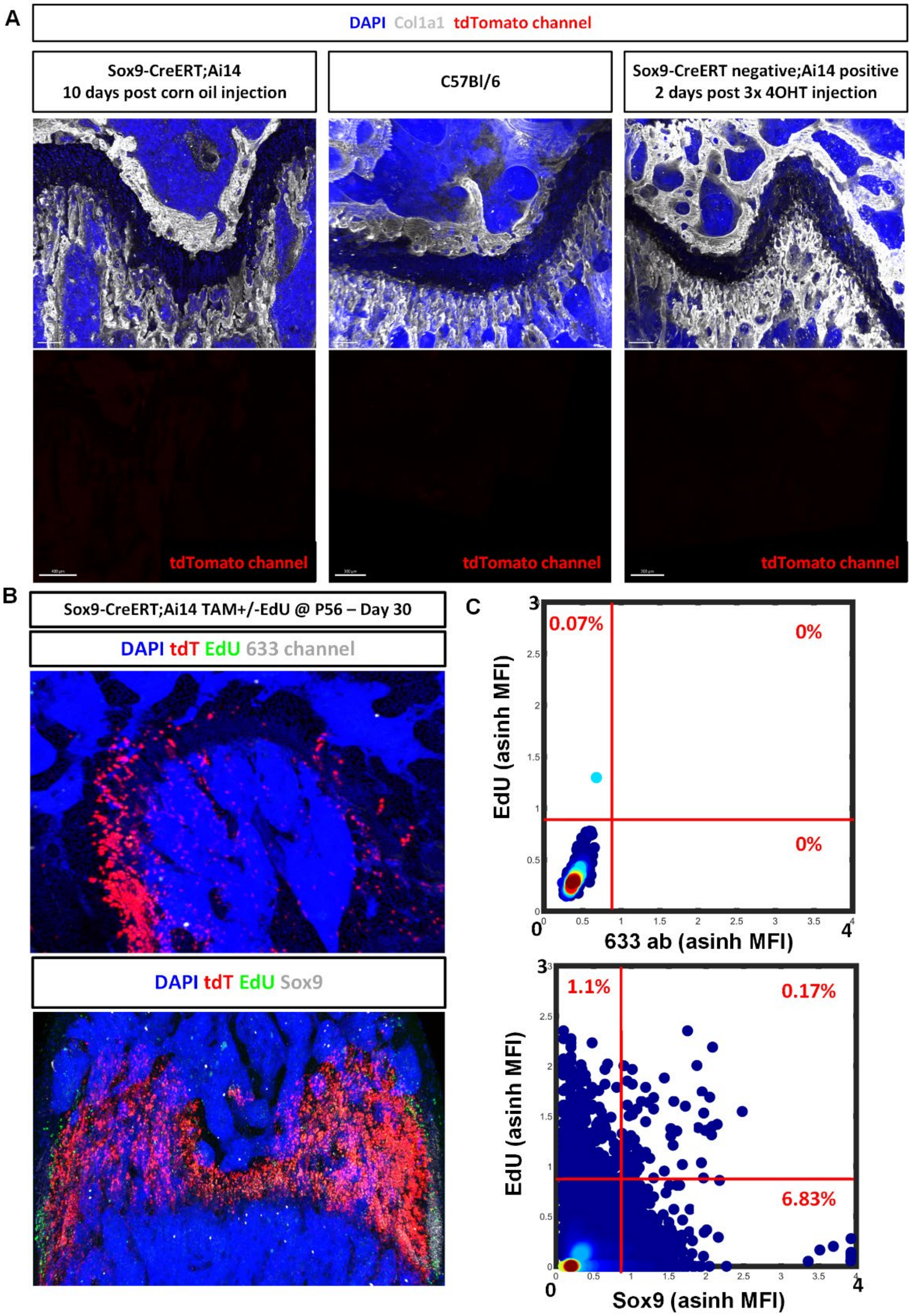


**Figure S1. Staining, gating and imaging controls.** A) Sox9-CreERT;Ai14 were pulsed with corn oil and femurs were collected 10 days post injection. C57Bl/6 mice femurs were collected without any injections, and Sox9-CreERT positive;Ai14 negative mice were pulsed with three 4OHT injections and femurs were retrieved two days later. B) Sox9-CreERT;Ai14 were pulsed with three TAM +/- three EdU and femurs were retrieved 30 days later. All femurs collected were then processed by confocal microscopy and imaging cytometry. C) Gating strategy using controls.

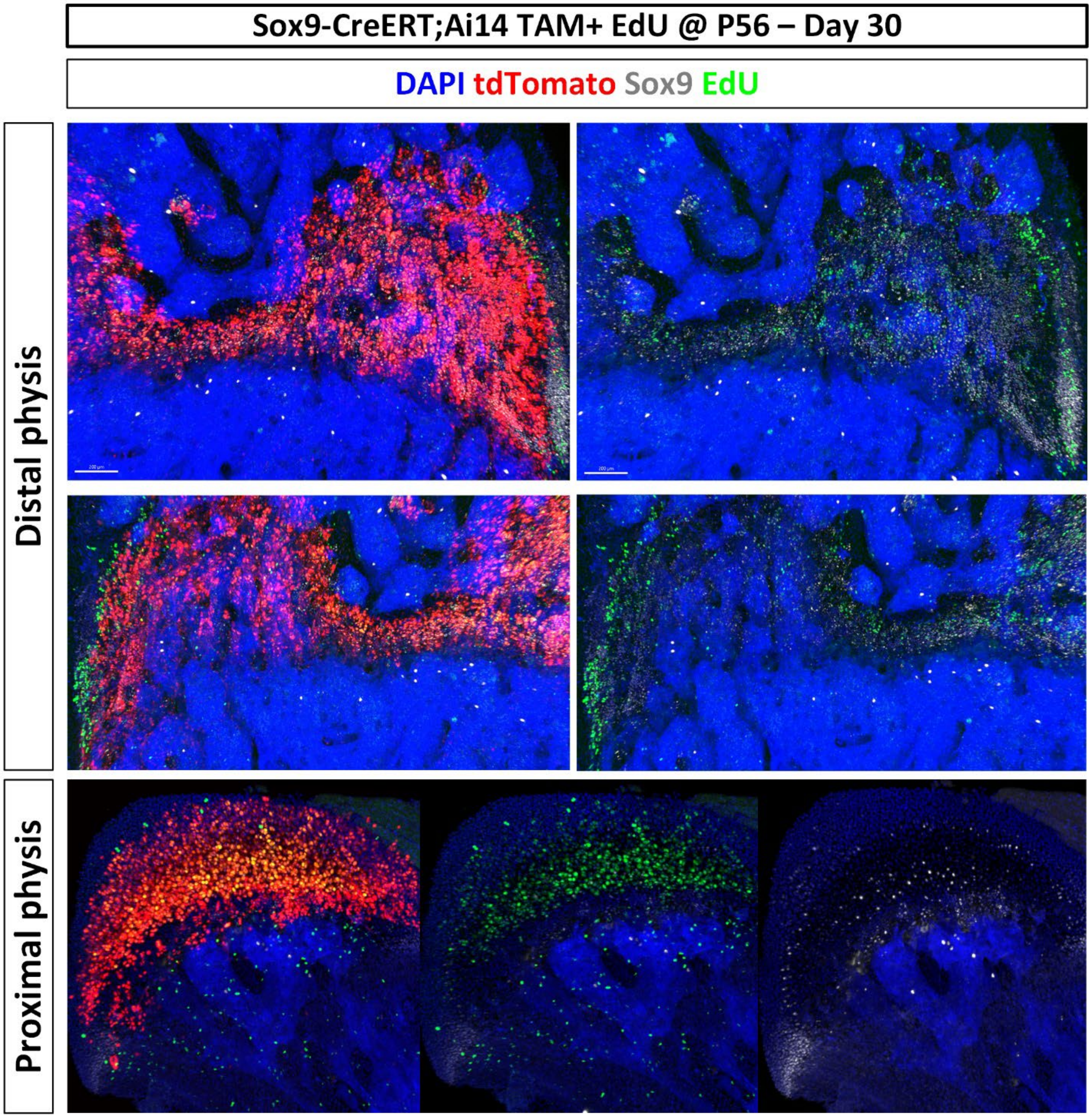


**Figure S2. Dual lineage tracing of Sox9+ cells in postnatal skeleton.** Sox9-CreERT;Ai14 mice received TAM and EdU on three consecutive days (P54-P56), femurs were retrieved on days 30 post-labeling to assess the fate of Sox9+ and proliferative cells. Samples were immunostained with anti-Sox9 antibody and counterstained with DAPI.

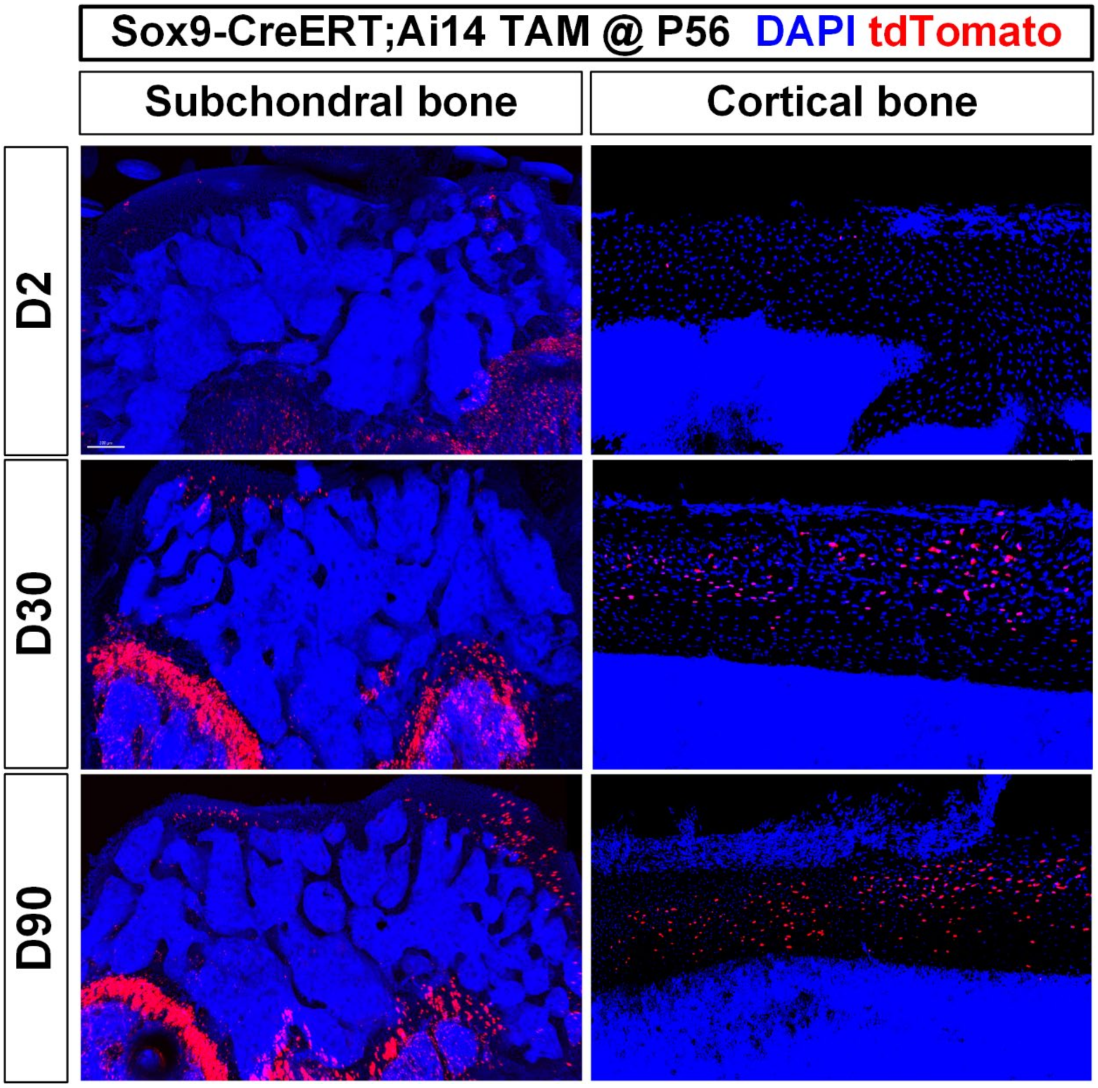


**Figure S3. Lineage tracing of Sox9+ cells in postnatal skeleton.** Sox9-CreERT;Ai14 mice received TAM injection at P56, femurs were then collected at D2, D30 and D90 post labeling, and processed for imaging. Samples counterstained with DAPI.

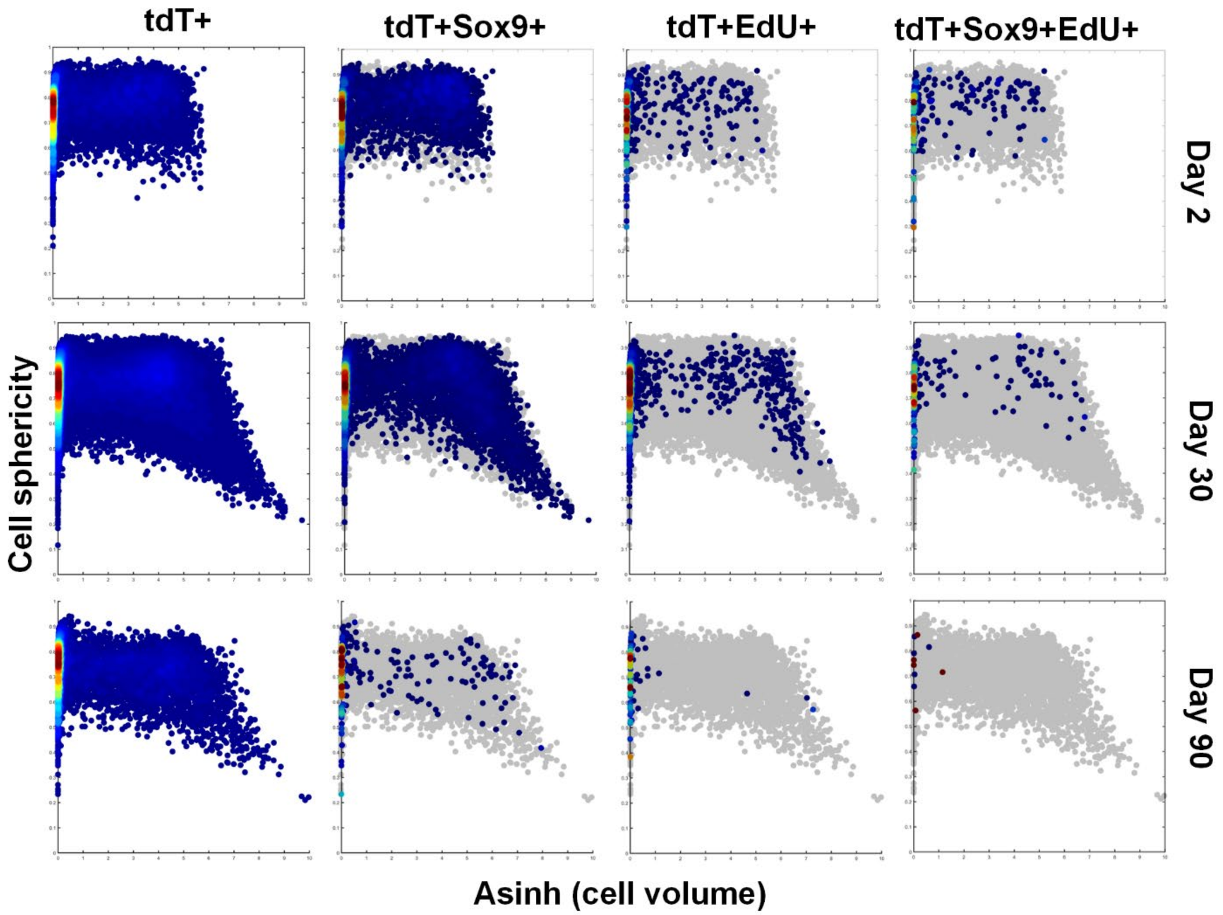


**Figure S4. Variation in cell shape and size during lineage tracing of Sox9+ mSSCs.** Lineage tracing data presented in Fig.1-2 displayed here to assess variation in cell shape (sphericity) and volume over time. Total segmented tdT+ cells (first column, density heatmaps, shown in grey in other columns) gradually become larger and less spherical as they differentiate into chondrocytes and osteocytes. However, cells that retain EdU and maintain Sox9 expression long term remain spherical and small.

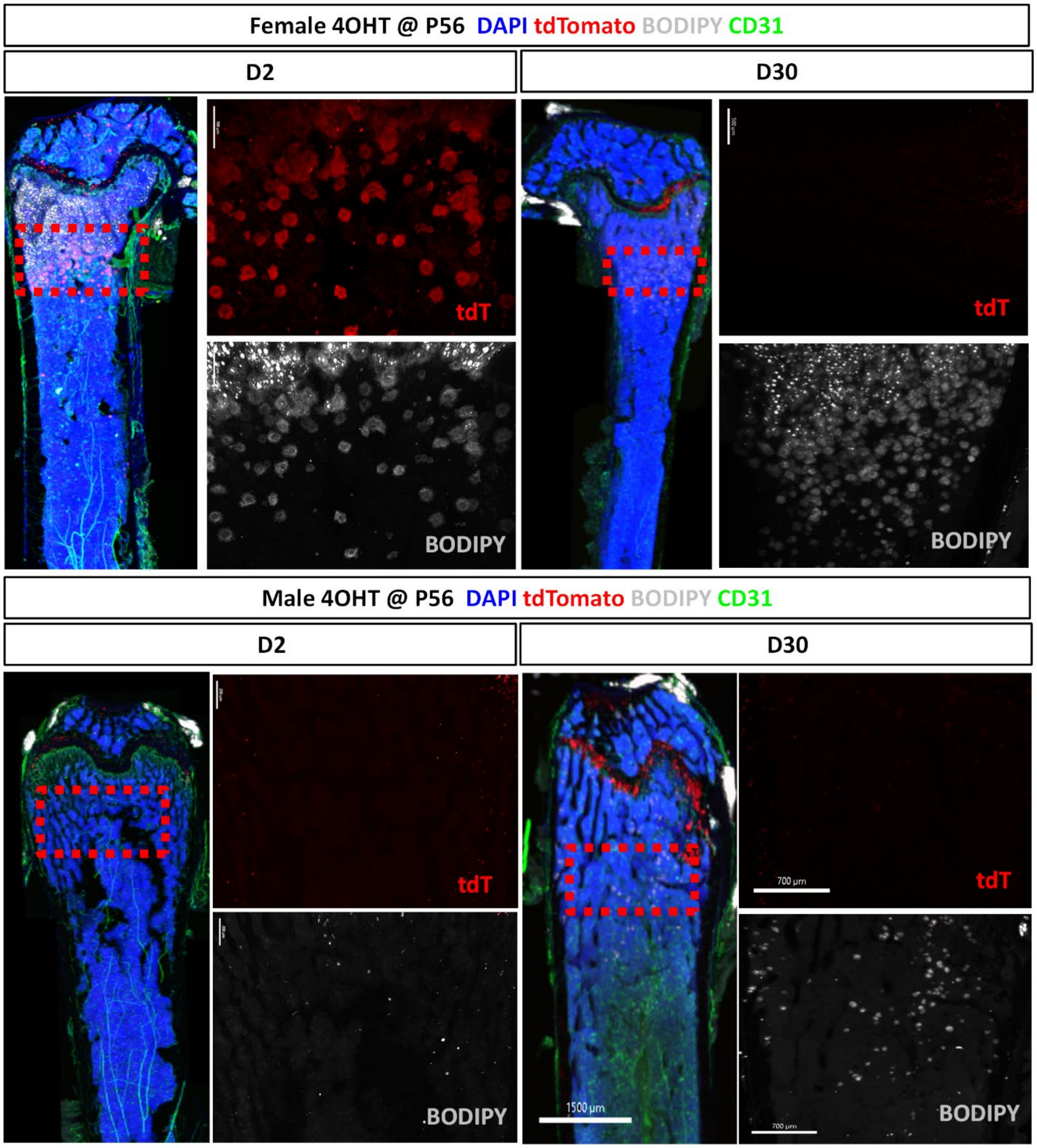


**Figure S5. Comparison of marrow adipocytes between males and females in juvenile mice.** Sox9-CreERT2;Ai14 mice were injected with 4OHT at P56, femurs were then collected at day 2 post injection and processed for immunostaining then analyzed by confocal imaging. BODIPY: lipid stain, CD31: endothelial cells marker.

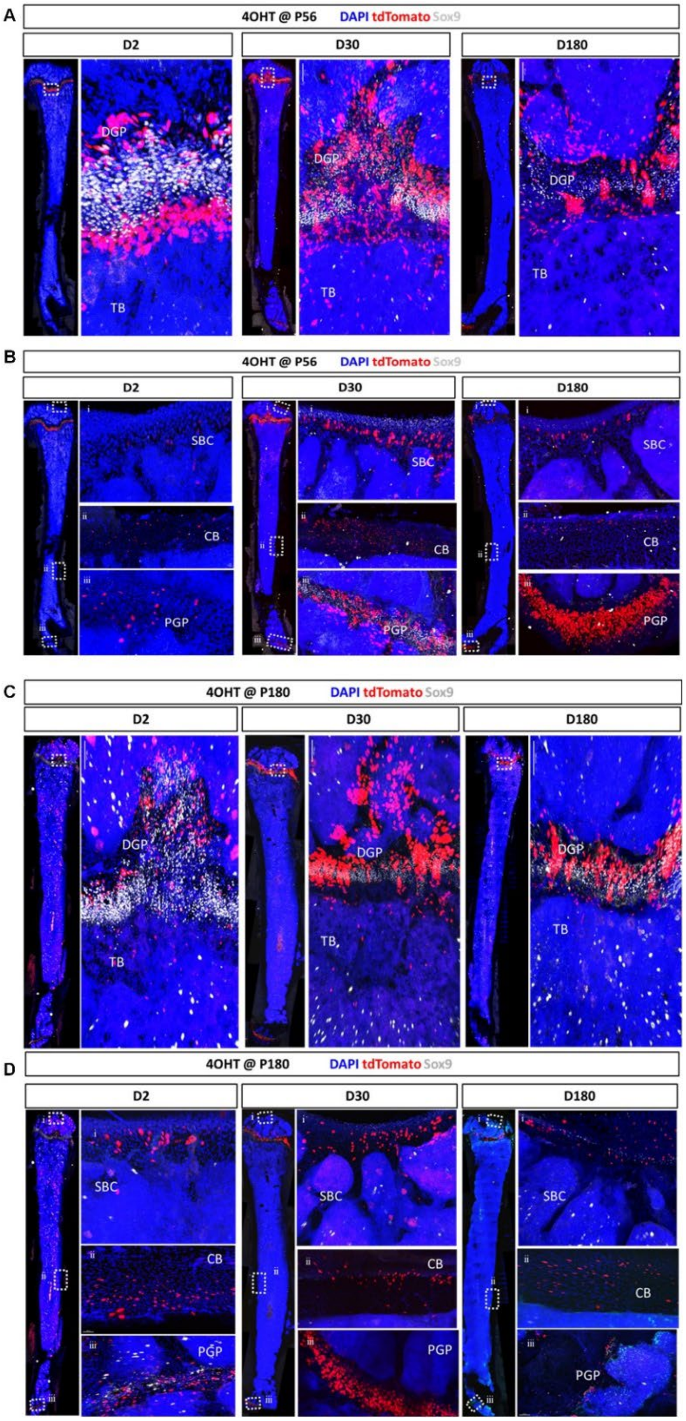


**Figure S6. Distribution of tdT+ and Sox9+ cells in juvenile and adult mice.** A, B) 4OHT was administered at P56 C, D) and at P180 in female mice inducing constitutive expression of tdTomato in Sox9-CreERT;Ai14 mice. tdT+ cells were chased at D2, D30 days and D180 post-4OHT injection, then immunostained and analyzed by confocal microscopy. SBC: Subchondral bone; CB: Cortical bone; PGP: Proximal growth plate; DGP: Distal growth plate; TB: Trabecular bone.

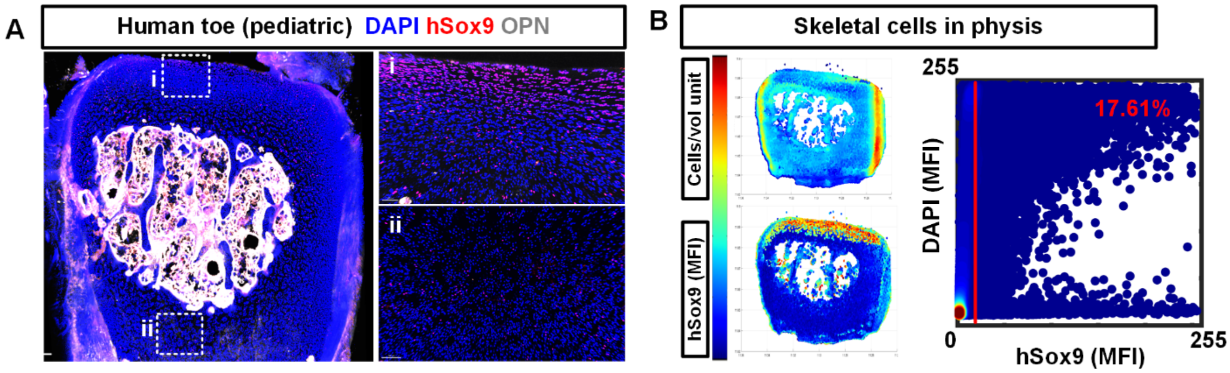


**Figure S7. Sox9+ putative osteochondral SSCs in human postnatal skeletal tissue.** A) Phalanx of a toe obtained from a pediatric patient with polydactyly whose toe was surgically amputated. The bone was processed similarly to mouse bones, immunostained for hSox9 and osteopontin (OPN), and counterstained with DAPI. The hSox9 antibody shown here is different from the one shown in Figure 6. B) Quantification of hSox9 in cells located in the physis of the bone shown in A.

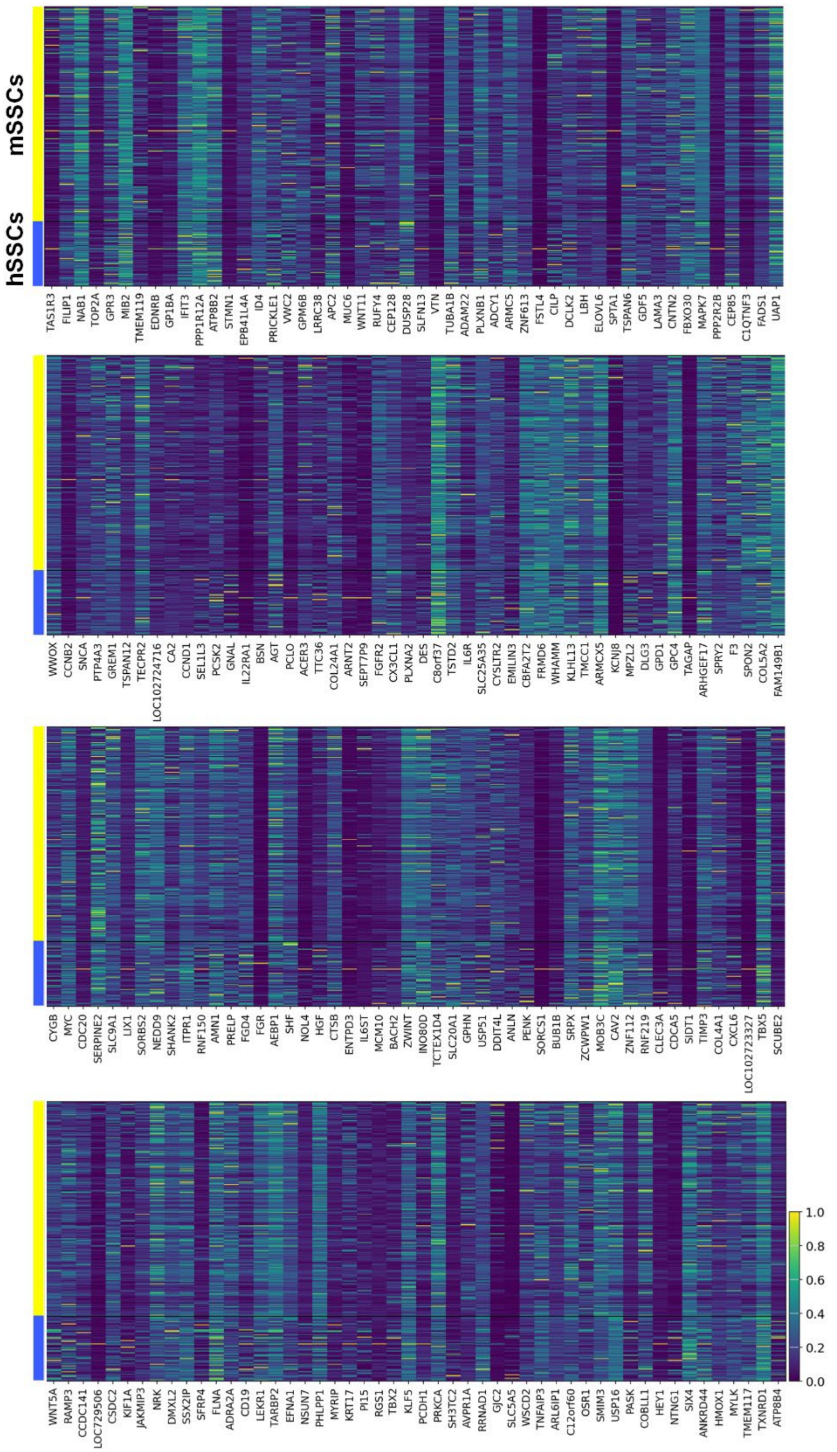


**Figure S8. Gene expression profile comparison between mSSCs and hSSCs.** Top 200 genes that show conserved expression in both mSSCs and hSSCs.

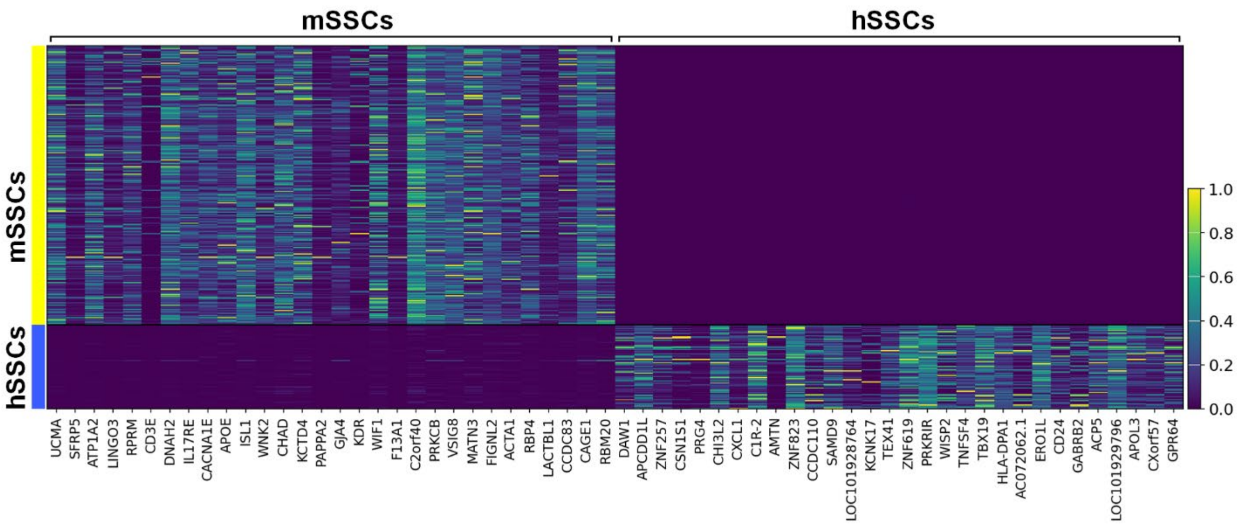


**Figure S9. Gene expression profile comparison between mSSCs and hSSCs.** Top 30 genes that the most differentially expressed between mSSCs and hSSCs.

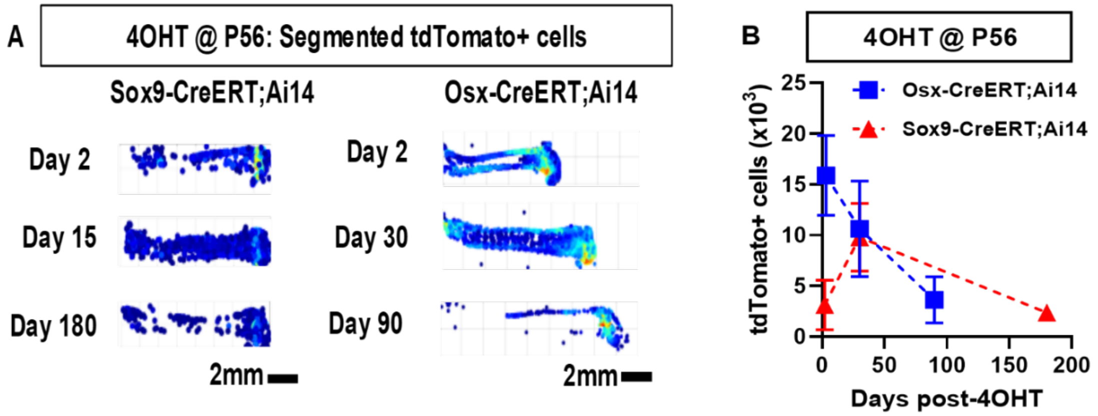


**Figure S10. Comparison of lineage tracing of skeletal cells using two different Cre-driver lines.** A) 4OHT was administered at P56 in female mice, inducing constitutive expression of tdTomato in Osx-CreERT+ (left) or Sox9-CreERT+ (right) cells and their progeny upon cell division. Shown are heatmaps of segmented tdTomato+ cells' position and density. B) Quantification of segmented tdT+ cells over time (n=3 for each data point).

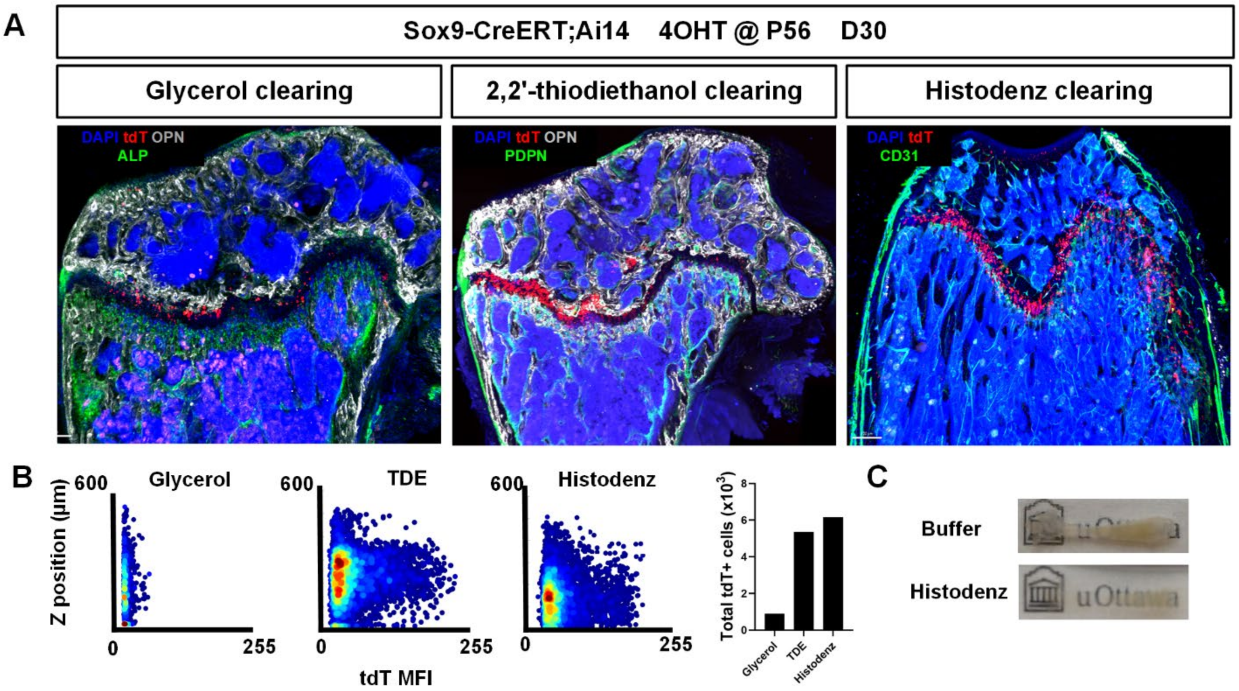


**Figure S11. Comparison of different clearing solution.** A) Sox9-CreERT;Ai14 mice received 4OHT injection at P56, femurs were then collected at D30 post injection and processed by confocal microscopy and imaging cytometry. B) Xit plots showing the depth of imaging (position) versus mean fluorescence intensity (MFI) of tdT for each clearing solution. C) Thick section of a mouse femur before and after optical clearing with Histodenz.

**Table 1. Antibodies used**

| Catalog number | Target | Alternative names | Clone | Company | Target Species | Host Species |
|---|---|---|---|---|---|---|
| AF808 | OPN | osteopontin, SPP1, bone sialoprotein 1 | poly | R&D Systems | mouse | goat |
| CL50151AP | collagen type 1 | col1a1 | poly | Cedarlane | mouse | rabbit |
| AF3628 | CD31 | PECAM1 | poly | R&D Systems | mouse | goat |
| AB5320 | NG2 | Melanoma chondroitin sulfate proteoglycan 3, Melanoma-associated chondroitin sulfate proteoglycan, chondroitin sulfate proteoglycan 4, CSPG4 | poly | Millipore/Sigma | Human, Mouse, Rat, Monkey | rabbit |
| ab185966 | Sox9 | | mono/EPR14335-78 | Abcam | Mouse, Rat, Human | rabbit |
| AF3075 | hSOX9 | | poly | R&D systems | human, mouse | goat |
| NBP1-85551 | hSOX9 | | poly | Novus Biologicals | Hu, Mu, Rt, Po, Ca | rabbit |
| AF3824-SP | ITGB5 | Integrin beta 5 | poly | R&D Systems | Human (50% mouse) | sheep |
